# Tracing active members in microbial communities by BONCAT and click chemistry-based enrichment of newly synthesised proteins

**DOI:** 10.1101/2024.08.27.609880

**Authors:** Patrick Hellwig, Daniel Kautzner, Robert Heyer, Anna Dittrich, Daniel Wibberg, Tobias Busche, Anika Winkler, Udo Reichl, Dirk Benndorf

## Abstract

A comprehensive understanding of microbial community dynamics is fundamental to the advancement of environmental microbiology, human health, and biotechnology. Metaproteomics, i.e. the analysis of all proteins in a microbial community, provides insights into these complex systems. Microbial adaptation and activity depend to an important extent on newly synthesized proteins (nP), however, the distinction between nP and bulk proteins is challenging. The application of bioorthogonal non-canonical amino acid tagging (BONCAT) with click chemistry has demonstrated efficacy in the enrichment of nP in pure cultures. However, the transfer of this technique to microbial communities has proven challenging and has therefore not been used on microbial communities before. To address this, a new workflow with efficient and specific nP enrichment was developed using a laboratory-scale mixture of labelled *E. coli* and unlabelled yeast. This workflow was successfully applied to an anaerobic microbial community with initially low BONCAT efficiency. A substrate shift from glucose to ethanol selectively enriched nP with minimal background. The identification of bifunctional alcohol dehydrogenase and a syntrophic interaction between an ethanol-utilizing bacterium and two methanogens (hydrogenotrophic and acetoclastic) demonstrates the potential of metaproteomics targeting nP to trace microbial activity in complex microbial communities.

## Graphical Abstract

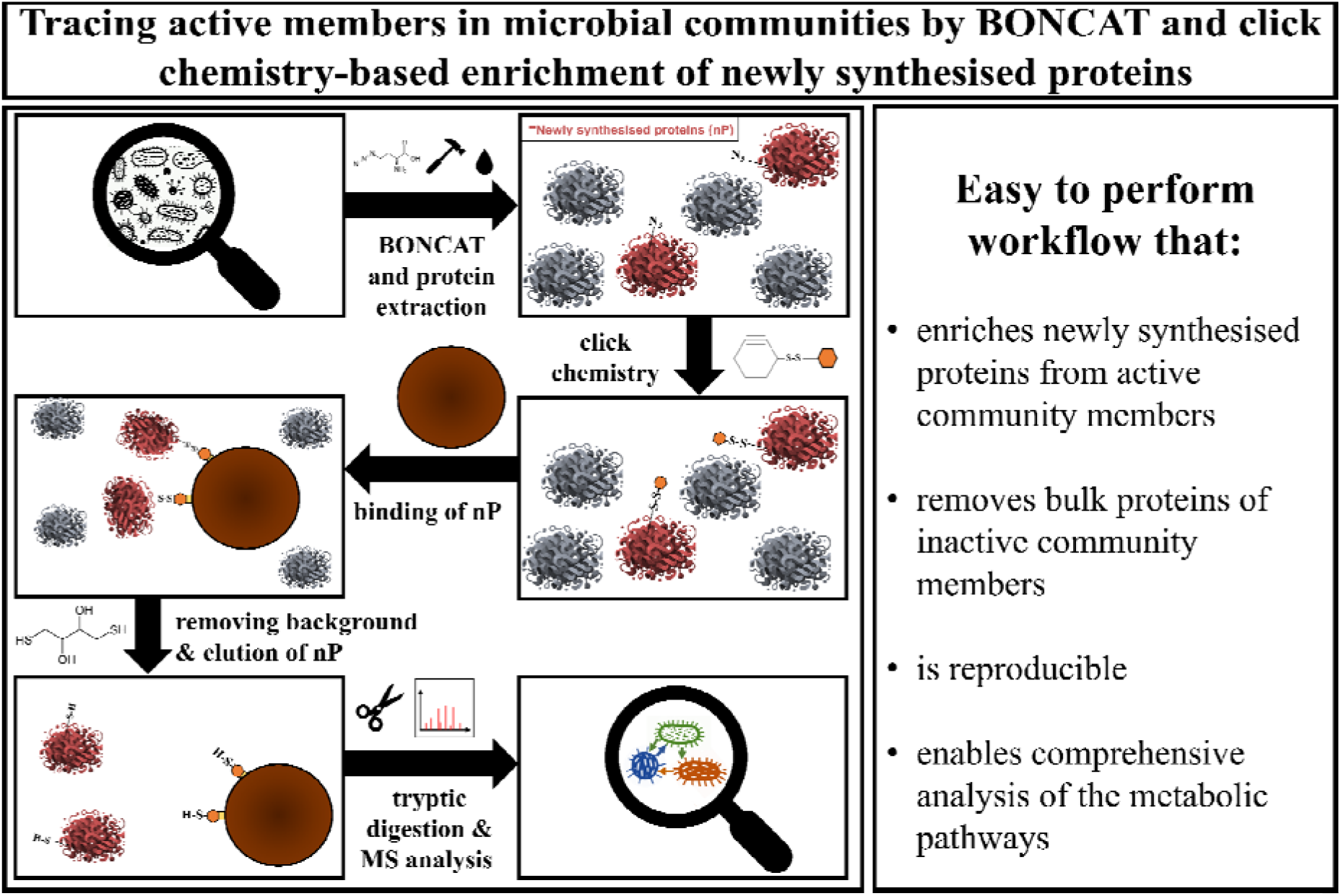

## Introduction

Microbial communities (MC) include single-celled eukaryotes, archaea, bacteria, and viruses [1–3]. These communities are found in various environments, including soil and water, and within living organisms such as animals and humans. Additionally, MC are used for industrial processes such as wastewater treatment or energy production e.g. in anaerobic digestion (AD) [4]. The understanding of MC is still very limited due to their inherent complexity. Enhancing our understanding will help to cure diseases, optimize biotechnological processes, and even act against climate change [5–7]. Due to its immense importance in the analysis and characterization of MC gained prominence in the last decades.

Metagenomics is currently the most widely used method to characterise MCs. It involves the simultaneous genomic analysis of all members of an MC and provides insight into the taxonomic composition and metabolic potential of an MC [8]. However, only in combination with other meta-omics techniques such as metaproteomics, which analyses all proteins of an MC, it is possible to fully decipher the functions of the MC [9].

Microbial adaptation to changing environments or process parameters and the resulting changed microbial activity occurs primarily through the expression of newly synthesized proteins (nP), which can be hardly distinguished from already present bulk proteins. Consequently, the analysis of the nP would provide a novel dimension of metaproteomics, thereby facilitating a deeper comprehension of MC activities.

The standard method of tracking nP is stable isotope labeling (SIP). In protein-SIP stable isotopes, such as ^13^C, ^15^N, and ^2^H (deuterium), are introduced into the cultivation media. These isotopes are incorporated into amino acids and the modified amino acids will be incorporated in nP [10, 11]. This allows a distinction between labeled nP from non-labeled proteins. Alternatively, advances in bioorthogonal chemistry have enabled another way of identification of nP. One technique is bioorthogonal non-canonical amino acid tagging (BONCAT), which allows the selective labelling of nP [12]. In BONCAT, analog non-canonical amino acids (ncAA) are added to cells or MCs. During protein synthesis, the tRNA-amino acyl synthetase of an amino acid mistakenly selects the ncAA instead of the canonical amino acid. 4-Azido-L-homoalanine (AHA) and L-Homopropargylglycine (HPG). Both are the ncAAs replacing methionine [13]. Note, incorporation of AHA and HPG does not depend on genetic modification of tRNAs in the uptaking cells. Studies in *E. coli* showed that the likelihood of incorporation is 1:390 for AHA and 1:500 for HPG [13]. The incorporation rate of ncAA into the amino acid chain of a protein correlates negatively with the concentration of the targeted amino acid in the medium [14]. This amino acid should therefore be avoided. However, Ignacio et al. (2023) [15] have recently developed a method called THRONCAT, in which ncAA incorporation is not affected by external canonical amino acid concentrations, which could significantly expand the range of applications for ncAA labeling in the future. The incorporation of ncAAs into nP results in a mass shift that can be detected by mass spectrometry (MS). Nevertheless, the low proportion of labelled nP in relation to the bulk proteins in MC is so far insufficient to permit the reliable identification of nP through metaproteomics.

As a remedy, BONCAT can be combined with click chemistry (CC) in the form of azide-alkyne cycloaddition [16, 17]. Thus, a fluorescence or biotin marker can be attached to the AHA (azide group) or HPG (alkyne group) [18]. BONCAT and CC have previously been applied to visualize and enrich nP from eukaryotic cells, such as neurons [14], HeLa cells [19], or HEK cells [20] and from viruses and their host cells [21]. However, BONCAT combined with CC has been used mainly in cell culture experiments with only one cell type. Hatzenpichler et al. [22] showed that the application of BONCAT in MC is possible and discussed the advantages and disadvantages of BONCAT and its application to MC in great detail [23]. Additionally, Reichart et al. [24] showed that sorting active ncAA-labelled microbes from non-active microbes with flow cytometry is possible. BONCAT and CC also show promise for bacteriophages in MC [25, 26].

In the case of metaproteomics, the detection of nP is of paramount importance for the understanding of MC. However, the currently available enrichment methods for nP are designed for pure cultures with high ncAA-labeling. In contrast, low enrichment efficiencies of nP and an enormous, unspecific binding of bulk proteins are achieved when using more complex and less ncAA-labelled MC samples. Therefore, there is a clear need for the development of reliable, specific, and reproducible methods for the detection of nP in MC *via* MS. The aim of this work was thus to develop a workflow that allows the enrichment of nP from growing MC for the detection *via* MS. The main focus was on the enrichment of AHA-labelled nP since the conditions of BONCAT have to be individually adapted to each experimental condition and MC in particular to the dynamics of protein translation [22, 27]. Therefore, we developed a test system for nP enrichment. It should demonstrate the specificity of nP enrichment and be easy to reproduce in any laboratory. Based on these restrictions, a test system consisting of *E. coli* and *Saccharomyces cerevisiae* (yeast) was chosen. *E. coli* was labelled with AHA after a substrate shift from glucose to lactose that induces metabolic changes and thus expression of nP involved in lactose metabolism. Next, unlabelled yeast cells were added. The established workflow allowed the enrichment of *E. coli* nPs involved in lactose metabolism without contamination by yeast proteins.

As a proof of concept, the workflow was applied to enrich nP from an anaerobic MC derived from a laboratory-scale biogas reactor (LBR). Given the slow growth of this anaerobic MC, the efficiency of ncAA-labeling was expected to be suboptimal. However, if nP can be enriched from this MC and reliably detected in MS, the developed workflow should be applicable to other slowly growing MC. The LBR used for this study was operated with glucose as the primary substrate. The objective was to shift the substrate to ethanol so that the nP associated with the consumption of ethanol as a secondary substrate would be labelled with AHA and could subsequently be enriched with the established workflow. It was anticipated that successful nP enrichment would allow to trace the degradation pathway of ethanol under anaerobic conditions. It may even be possible to gain new insights into this poorly described metabolic pathway [28].

## Material and Methods

In the following we briefly describe the workflow established (Figure 1). A detailed description of the methods, including a step-by-step standard operation procedure (SOP), can be found in Supplementary Note 1. All cultivation experiments were performed in triplicates.

**Figure 1:**
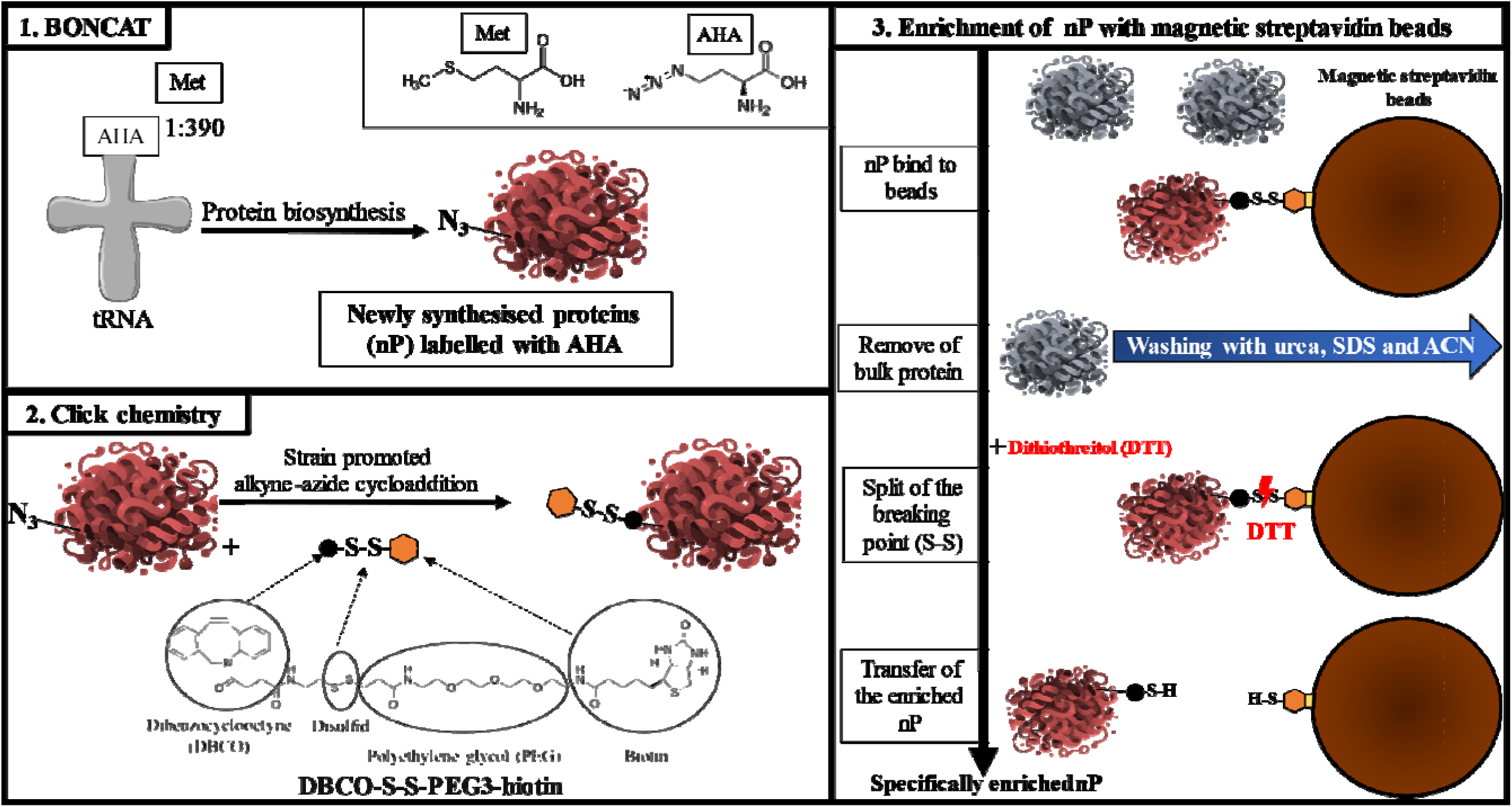
Workflow of the enrichment of newly synthesized proteins. **1.:** nP were labelled with AHA. **2.:** Free cysteines of the proteins were blocked with IAA (not shown). The labelled nP were tagged with DBCO-S-S-PEG_3_-biotin within 1 h. **3.:** The biotin-tagged nP were bound to magnetic streptavidin beads (blocked with amino acids in advance). All proteins not bound to the beads were removed by harsh washing with urea, sodium dodecyl sulfate (SDS), and acetonitrile (ACN). The bound proteins were eluted from the streptavidin beads with DTT. The specifically enriched nP were transferred to a new tube, tryptic digested, and further analysed with LC-MS/MS.

### AHA-labeling of *E. coli* nP

The carbon source of an E. coli (DSM 5911) culture grown aerobically at 37°C in M9 media was switched from glucose to lactose while the cells were incubated with 100 µM AHA (sterile filtrated). The same substrate shift was done for control cells without adding AHA. After the harvest, the two *E. coli* cultures were separately mixed with an overnight yeast culture. The mixture of *E. coli* and yeast is named “test system” in the following (see SOP 1).

### AHA-labeling of nP from MC of LBR

50 mL of cell suspensions from an LBR MC (1 L total volume, 40°C, 75 rpm, pH 7.4 – 7.5, Supplementary Table 1) was sampled using a 50 mL syringe. The cell suspensions were then transferred to nitrogen-flushed 100 mL serum bottles, diluted 1:1 with 50 mL anaerobic medium without glucose (Supplementary Table 2), and incubated for one hour. Following this, 1 mL of 10% (vol./vol.) ethanol solution was added. Additionally, for BONCAT, 1 mL of 20 mM AHA was introduced, while controls received 1 mL of water. An extra control was prepared with 3.8 mM glucose instead of ethanol. The cultures were sealed and incubated for 24 hours at 40°C with gentle stirring until harvest.

### Cell disruption with a ball mill

Samples from the test system and the LBR MC were suspended in 5 mL of 20 mM Tris/HCl- MgCl buffer (pH 7.5) and treated with 1 g/mL of silica beads and 0.5 µL/mL of Cyanase^TM^ Nuclease solution. The cells were lysed in a ball mill, incubated at 37°C for 15 minutes, and then centrifuged. The resulting supernatant was transferred to a new reaction tube(see SOP 1).

### Protein extraction with acetone (for pure cultures)

The proteins of the test system were precipitated with the five-fold volume of ice-cold acetone at -20°C for at least 1 h followed by centrifugation. The supernatant was discarded, and the protein pellets were dried under a fume hood at room temperature (RT). The dry protein pellets were resuspended in incubation buffer I. The protein samples were centrifuged, and the supernatants were transferred to new reaction tubes (see SOP 1).

### Protein extraction with methanol and chloroform (for environmental samples)

The MC proteins were extracted using a mixture of methanol, chloroform, and water, followed by centrifugation [29]. After carefully removing the top phase, methanol was added, and the mixture was vortexed and centrifuged again. The resulting protein pellets were dried and resuspended in incubation buffer II. After centrifugation, the supernatant was transferred to new tubes.

### Protein quantification

Amido black assay was used to quantify the protein concentration of each sample, as described earlier [30].

### Click chemistry with fluorophores and SDS-PAGE with fluorescent scan

AHA incorporation was assessed using 10 μg protein from each sample. For this purpose, the dye Dibenzocyclooctyne (DBCO) Cyanin 5.5 was attached using CC following SOP 2 in Supplementary Note 1. The fluorophore-tagged proteins were separated in a 1.5 mm 10 % SDS-PAGE and scanned twice: once with a LI-COR Odyssey Classic fluorescence scanner at 700 nm and a second time after Coomassie staining [30, 31]. (see SOP 5).

### Click chemistry with biotin linker

Based on the fluorescence signal (previous paragraph), 100 μg protein of the test system and 200 μg protein of the MC were used for the attachment of DBCO-SS-Biotin via CC (see SOP 2).

### Enrichment of AHA-labelled proteins

The nP of the test system and the MC were enriched using Dynabeads™ MyOne™ Streptavidin C1 beads and the protocol described in SOP 3. The beads were blocked with bovine serum albumin (BSA), amino acids (AA), or left blocked.

### FASP digestion

The tryptic digestion was performed using FASP [30] with few modifications (see SOP 4). The proteins contained in the dithiothreitol (DTT) elution phase and in the elution phase of the SDS boiled beads were precipitated with acetone. 25 µg protein of each sample before CC (“not enriched samples”) was precipitated with acetone (control without enrichment). The resulting protein pellets were dissolved in 200 μL 8 M urea buffer and transferred to a filter unit (Centrifugal Filter Unit, 10 kDa). In FASP digestion, a ratio of 1:100 (trypsin quantity to protein quantity) was used. For the enriched nP fractions, 100 ng trypsin was used. After extraction and concentration via vacuum centrifuge, the eluted peptides were transferred to HPLC vials for MS measurement.

### LC-MS/MS measurements

The peptides of each sample were measured using a timsTOF™ pro mass spectrometer (Bruker Daltonik GmbH, Bremen, Germany) coupled online to an UltiMate® 3000 nano splitless reversed-phase nanoHPLC (Thermo Fisher Scientific, Dreieich, Germany) in PASEF® mode.

### Illumina library preparation, MiSeq sequencing, and metagenome assembly for protein database

Three samples of the LBR used in this study were collected in different weeks for metagenomic sequencing to generate a protein database for the metaproteomic analysis in this study. The samples from the LBR were sequenced independently using PCR-free libraries prepared with the Illumina TruSeq® DNA PCR-free kit [32]. The libraries were subjected to quality control and sequenced on the MiSeq platform (2 × 300 bp paired-end, v3 chemistry). After sequencing, data were stripped of adapters and low-quality reads [33], followed by assembly with Megahit (v1.1.1) [34] and gene prediction with Prodigal v.2.6.0 [35]. Metagenomic binning [36, 37] was performed with Bowtie 2 [38] and MetaBAT2 (76 Metagenome assembled genome (MAGs)) [39], while completeness and contamination were assessed with BUSCO (v5.7.0) [40]). The translated amino acid sequences of the predicted genes were generated and the replicates were merged for a protein database generation with Contig informations. For the most abundant MAGs, the MAG sequence was uploaded to the Type (Strain) Genome Server (TYGS) to evaluate the taxonomy [41, 42].

### Protein identification using Mascot and the MetaProteomeAnalyzer

With Compass Data Analysis software (version 5.1.0.177, Bruker Corporation, Bremen, Germany) the raw files were converted into Mascot Generic Files (mgf) and mgf were searched with Mascot (version 2.6) for the peptide spectrum matches. For the test system, a defined UniProtKB/SwissProt database (08/05/2022) containing *E. coli* K12 (taxonomy_id:83333) and yeast (taxonomy_id:4932) proteins was used. A metagenome from the LBR was used for the MC samples (see Illumina library preparation, MiSeq sequencing, and metagenome assembly). Metaproteins were generated from the MC samples using MetaProteomeAnalyzer software (version 3.0 [43]).

### Further analysis of the protein data

The protein data were uploaded to Prophane (version 6.2.6) [44] for basic local alignment search tool analysis of the functional and taxonomic annotation. For all unknown KO numbers after Functional Ontology Assignments for Metagenomes annotation [45], KofamKOALA [46] was used to identify possible missing KO numbers (Supplementary Table 3 and Supplementary Note 1).

## Results and Discussion

### Development of a nP enrichment workflow with the test system

Prior to developing the enrichment protocol, the *E. coli* (AHA-labelled) and yeast (unlabelled) test system was refined by optimising the AHA labelling of *E. coli,* measured by detecting the proportion of fluorescent *E. coli* cells generated through click chemistry. (Supplementary Figure 1). Attempts were made to enrich nP of *E. coli* from the test system with known enrichment protocols (e.g., [20, 21, 47]) but these were unsuccessful (data not shown). The major issues were that MS of tryptic digests was either unable to detect any proteins, or detected similar proportions of yeast proteins (unspecific background) and *E. coli* proteins. The results highlighted the need for a test system to monitor nP enrichment, and the importance of implementing negative and positive controls to regulate non-specific binding in each experimental procedure.

A first successful enrichment of nP after click chemistry was obtained with DBCO-SS-PEG3- biotin and the use of Dynabeads™ MyOne™ Streptavidin C1 beads (Supplementary Figure 2). The nP were eluted under mild conditions from streptavidin beads by the addition of DTT through the reductive cleavage of the disulfide bridge in the DBCO-SS-PEG3-biotin. Therefore, DBCO-SS-PEG3-biotin appeared to be an optimal choice for our nP enrichment. However, many unspecific yeast proteins (background) were co-eluted with the nP (Supplementary Figure 2).

Therefore, washing was optimized. An adapted washing strategy [20] and the use of phosphate-buffered saline (PBS) buffer with increased NaCl concentration (237 mM) reduced the amount of background yeast proteins. However, it strongly reduced the reproducibility of nP elution (Figure 2 A). For higher reproducibility of enrichment, the loss of nP during elution should be minimized.

**Figure 2:**
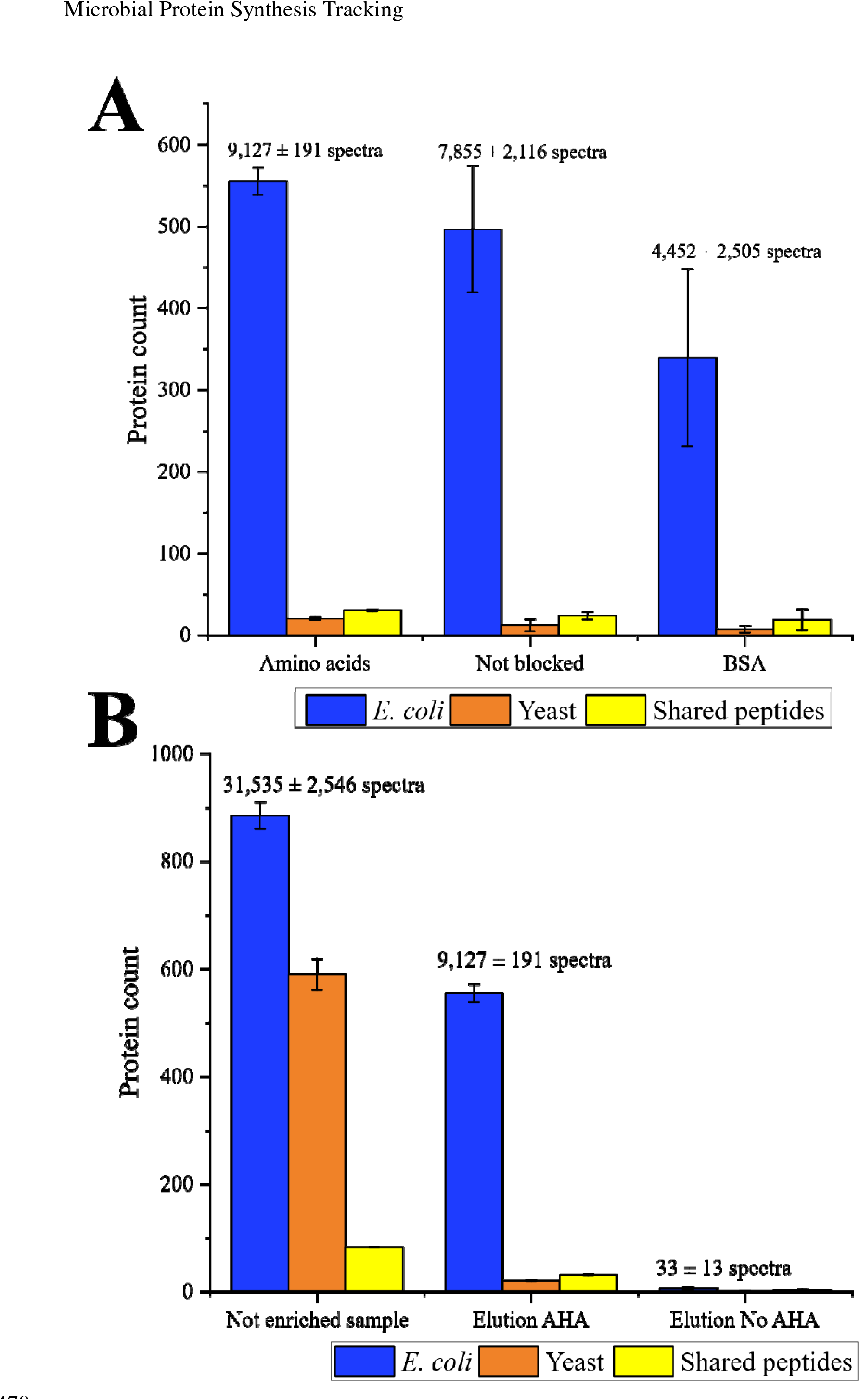
Number of identified proteins of the test system using the new enrichment method for nP. **A:** Comparison of the number of identified proteins from the test system between the tested methods for nP enrichment. All proteins identified in the elution with AHA were compared between the three enrichment methods: blocking with amino acids, blocking with BSA, and no blocking. **B:** The different fractions of nP enrichment with the AA method were compared. The number of proteins identified in the not enriched and AHA labelled sample (“Not enriched sample”), in the elution fraction of AHA labelled sample (“Elution AHA”), and in the elution fraction without AHA labelling (“Elution No AHA”) are shown. All proteins with at least two spectra were considered. The proteins were grouped into the taxonomies *E. coli* and yeast. Proteins with shared peptides were grouped as "Shared peptides". The bars represent the average of each group, and the error bars represent the standard derivation of three technical replicates. The raw data can be found in Supplementary Table 2.

Blocking the beads with 1 mg/mL BSA (e.g. [48, 49]), before loading the samples, reduced the amount of yeast protein background further but unfortunately also increased the variability in nP elution (Figure 2 A). Additionally, the elution phase was found to be contaminated with BSA proteins. Blocking the beads seemed to be important for the removal of non-specific protein, but BSA was not optimal.

Inspired by Nicora et al. (2013) [50], the beads were blocked with an AA blocking solution containing 16.67 µg/mL of leucine (aliphatic hydrophobic), tryptophan (aromatic hydrophobic), histidine (polar positive), glutamine (polar neutral), glutamic acid (polar negative) and glycine (nonpolar neutral) each. These AAs should not cause steric effects and should prevent any unspecific interactions with the beads. This method (“AA method”) maintained a low background and exhibited minimal variability and loss in nP elution (Figure 2 A), making it the most promising approach.

### Examining the AA method using the test system

In total, 1,558 ± 53 proteins (885 ± 25 *E. coli*, 590 ± 29 yeast) were identified in the AHA-labelled but non-enriched test system (“not enriched sample”) (Figure 2 B). Over a third of these proteins were yeast proteins. After nP enrichment, 607 ± 16 proteins (555 ± 17 *E. coli*, 21 ± 2 yeast) were identified, reducing the amount of identified yeast proteins by over 96% while retaining 63% of the *E. coli* proteins (Figure 2 B). Additionally, the AA method resulted in a minimal background in the unlabelled control test system (without AHA-labelled proteins) with 10 ± 3 proteins (6 ± 3 *E. coli*, 1 ± 1 yeast).

Furthermore, the AA method allowed the detection of 48 proteins that were absent in the not enriched samples (Supplementary Table 2). Of these, 29 were *E. coli* proteins that related to cell metabolism, movement, transport, and protein biosynthesis. Additionally, 19 very low-abundant yeast proteins were detected, likely co-eluting with AHA-labelled *E. coli* nP. This co-elution did not occur in the AA method control, which confirmed the specificity of the elution of labelled nP by DTT. Boiling of the beads with SDS-PAGE sample buffer released no additional proteins showing that DTT was very efficient in elution (Supplementary Figures 3 and 4). Consequently, the selected elution with DTT is not only specific but also highly efficient.

Due to the substrate shift from glucose to lactose, the enriched *E. coli* nP should include proteins responsible for lactose degradation encoded by the *lac operon,* namely β- galactosidase (*lacZ*), galactoside O-acetyltransferase (*lacA*), and β-galactoside permease (*lacY*) [51, 52]. Indeed, the AA method successfully enriched β-galactosidase and galactoside O-acetyltransferase by a relative foldchange of 3.7. However, β-galactoside permease was not detected, neither in the enriched sample nor in the not enriched sample. As it is an integral membrane protein it was probably lost during protein extraction [53].

Lactose consists of galactose and glucose. Therefore, the pathway for galactose metabolism proteins should also be enriched with the AA method. The assignment of the detected nP to the KEGG pathway for galactose metabolism showed that all enzymes for galactose degradation were identified and at least 2-fold enriched, including β-galactosidase (EC 3.2.1.23; 217 ± 6 spectra), aldose 1-epimerase (EC 5.1.3.3; 14 ± 1 spectra), galactokinase (EC 2.7.1.6; 50 ± 5 spectra), galactose-1-phosphate uridylyltransferase (EC 2. 7.7.12; 13 ± 3 spectra), UTP-glucose-1-phosphate uridylyltransferase (EC 2.7.7.9; 3 ± 1 spectra), and UDP-glucose 4-epimerase (EC 5.1.3.2; 34 ± 2 spectra) (Figure 3). Glycolytic enzymes were also identified; however, they did not exhibit a 2-fold increase in the enriched nP compared to the not enriched sample. This might be explained by the fact, that cells were already grown on glucose as substrate before the substrate shift (Figure 3). Furthermore, all dehydrogenases involved in the mixed acid fermentation of *E. coli* [54] were enriched after the substrate shift by the AA method(Supplementary Table 2). The substrate switch was carried out by centrifugation, which presumably led to anaerobic conditions for a short time and thus facilitated the formation of these enzymes. This further emphasizes the significance of nP as a conduit of knowledge regarding the activities of microorganisms.

**Figure 3:**
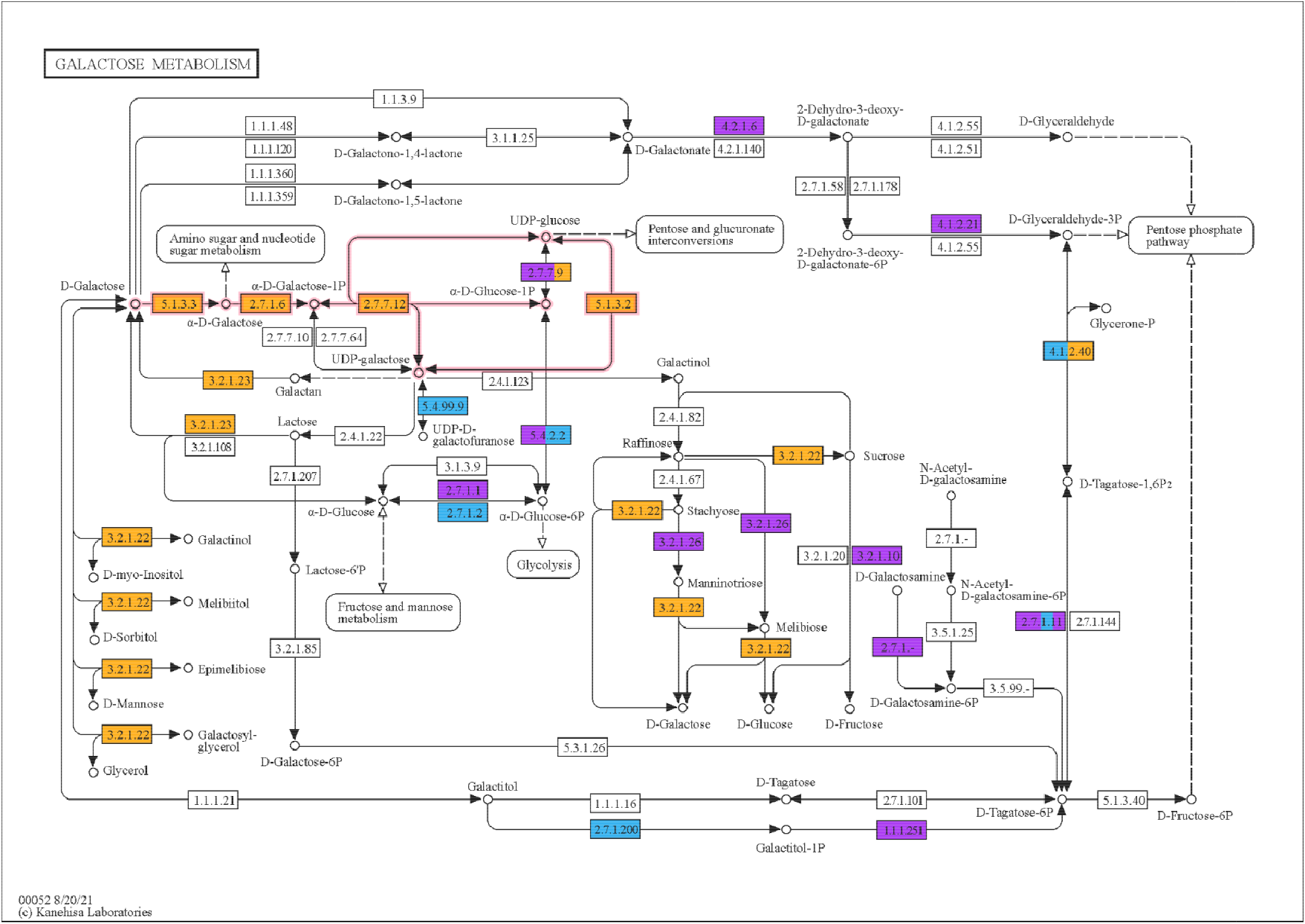
KEGG map of all identified proteins from galactose metabolism in *E. coli* in the test system (map00052). **Orange** indicates proteins enriched at least 2-fold with the developed AA method. **Blue** shows the proteins with no significant difference in abundance between the AA method and not enriched sample. **Purple** represents all proteins uniquely identified in the not enriched sample. Boxes with several colours show similar proteins with different abundances for this step. **Pink arrows** highlight the conversion pathway of galactose to α-D-glucose-1-phosphate. Blank boxes were not identified in any sample. Sample data were normalized by dividing the number of spectra of each protein by the sum of the spectra for each sample. The abundance of each protein was averaged over three replicates, and the relative abundance ratio between the not enriched sample and elution was calculated for comparison [98]

### Enrichment of nP in anaerobic MC of an LBR using the AA method

The next objective was to demonstrate that the established workflow is also effective for weaker AHA-labelled and more complex MC. Therefore, the nP of an anaerobic MC from an LBR were labelled with AHA while the substrate was switched from glucose to ethanol.

Initial quality control of successful AHA incorporation by SDS-PAGE revealed a weak fluorescent signal that represents nP tagged with DBCO-Cyanin 5.5 by CC (Supplementary Figure 5). The lower incorporation of AHA in the MC compared to a pure test sample reinforces the necessity for specific and effective enrichment of nP to reliably detect nP in MCs with MS. Unfortunately, in one out of three biological replicates of the experiment, no fluorescence was detectable (Supplementary Figure 5). Therefore, this replicate was excluded from further analysis. The other two replicates were processed with the AA method and subsequent metaproteome analysis.

To achieve more accurate identification of proteins within the LBR MC, a metagenome was sequenced from the LBR, and a corresponding protein database was generated. 76 MAGs were binned.

In total, 2063 (replicate 1) and 2065 (replicate 2) proteins were identified in the not enriched sample of the AHA-labelled MC (“Not enriched sample AHA”), while 525 (replicate 1) and 356 (replicate 2) proteins were identified after the enrichment of nP (“Elution AHA”).

Application of the AA method to the non-labelled control (“Elution no AHA”) showed a low non-specific protein background of 7 (replicate 1) and 25 (replicate 2) proteins (Figure 4 A). Notably, with the AA method, 26 proteins were identified (both replicates), which were absent in the not enriched sample (Figure 4 B). Consequently, the specific enrichment of nP should provide a better insight into the potential conversion of ethanol to methane in this LBR than the proteome analysed without enrichment.

**Figure 4:**
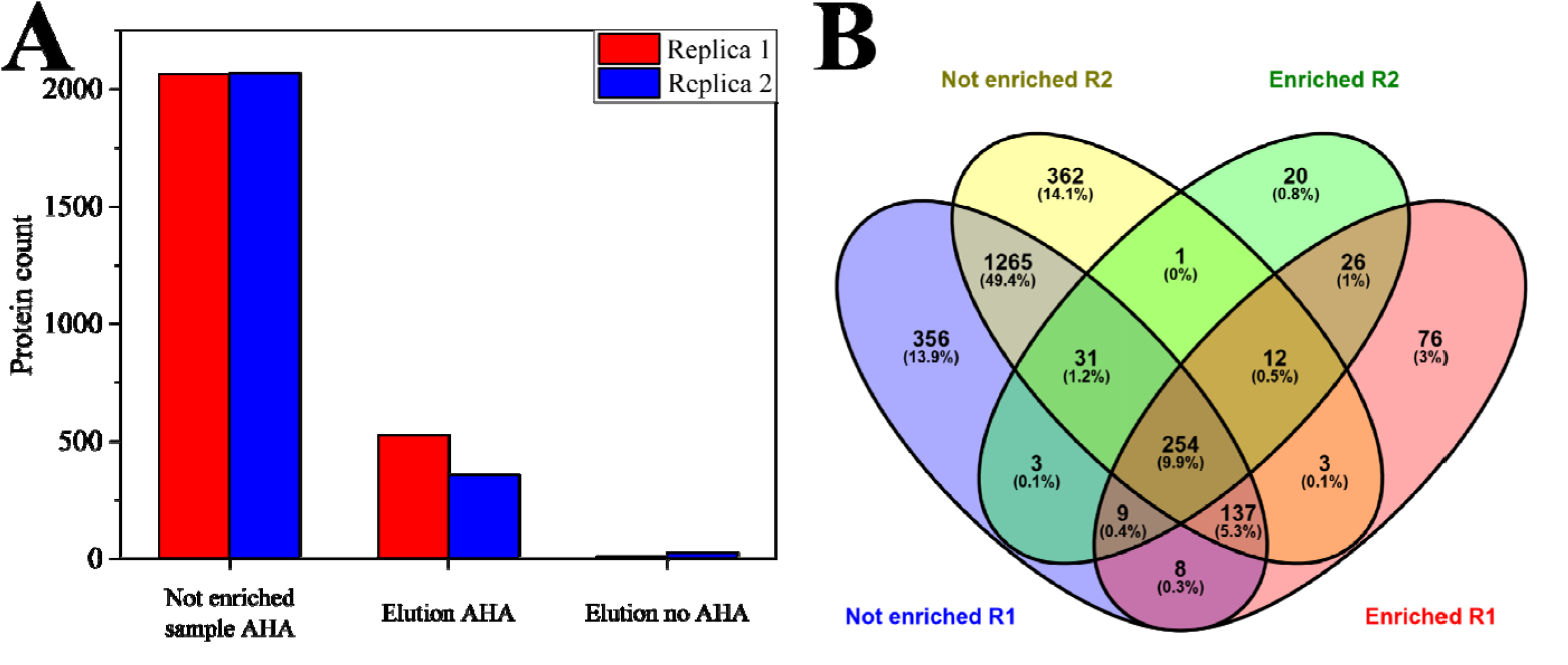
Number of identified metaproteins from AD using the AA method. The proteins were grouped based on shared peptides and the number of the resulting metaproteins was plotted. Each sample was prepared in biological duplicates, which are indicated in the plot by red and blue bars. The samples are the not enriched but AHA labelled sample (“Not enriched sample”), the enriched nPs of the AHA labelled sample in the elution fraction (“Elution AHA”), and in the elution fraction of the control without AHA labelling (“Elution No AHA”). **B:** The Venn diagram illustrates the complete set of proteins identified in the respective replicates of the non-enriched (“Not enriched”) and nP-enriched (“Enriched”) samples [99]. The raw data can be found in Supplementary Table 3.

### Ethanol-associated taxa based on nP enrichment

Initially, the enrichment of the nP was considered for the 20 most abundant MAGs of the AA method and without the AA method each based on their protein abundance divided by the MAG length (29 MAGs in total, Supplementary Table 6). The ratio of the MAGs between the AA method and without the AA method was calculated based on their protein abundance. The results of the two replicates demonstrate a consistent pattern of variation in protein abundance among the MAGs under investigation. Bin_27 (family *Propionibacteriaceae*), Bin_24 (class *Methanomicrobia*), Bin_41 (class *Clostridia*), and Bin 53 (species *Candidatus Methanoculleus thermohydrogenitrophicus*) are of particular interest due to their high protein abundance in all samples. The protein abundance of Bin_27 has been depleted by the AA method, whereas the proteins of the other three MAGs are similarly abundant in all samples. While certain MAGs, such as Bin_74 (family *Desulfobacteraceae*), Bin_62 (unknown *Bacteria*), Bin_46 (unknown *Bacteria*), Bin_11 (family *Synergistaceae*), and Bin_20 (family *Alcaligenaceae*), do not stand out due to their high protein abundance, they do exhibit strong protein enrichment with the AA method. In contrast, certain MAGs, including Bin_7 (family *Pseudomonadaceae*), Bin_23 (family *Methanobacteriaceae*), Bin_69 (family *Flavobacteriaceae*), Bin_71 (class *Clostridia*), Bin_75 (class *Actinomycetia*), and Bin_67 (family *Pseudomonadaceae*), exhibited a depletion pattern in their protein abundance after using the AA method (Figure 5).

**Figure 5:**
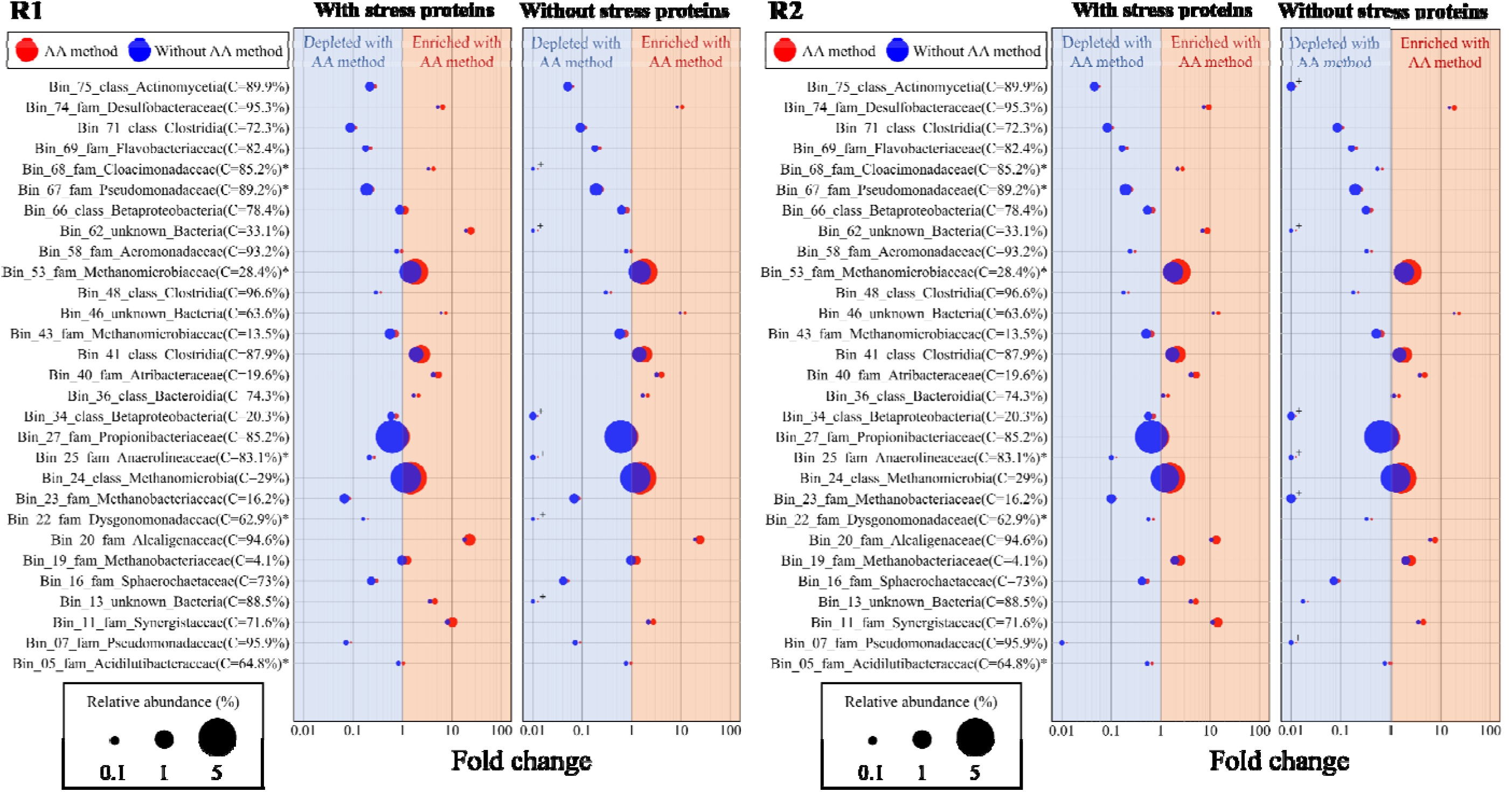
Bubble plot of the MAGs with the highest protein abundance identified using the AA method. The protein abundance of the 20 most abundant MAGs identified by the AA method and the 20 most abundant MAGs identified in the non-enriched sample were compared in 2 replicates (a total of 29 different MAGs). The fold changes were calculated based on the ratio of the relative spectral abundance of the MAG, which was normalised by the MAG length in the samples. The taxonomies were identified using the TYGS. The left bubble plot of each replicate shows the MAG abundance based on all identified proteins and the right without stress proteins. “C” in brackets shows the completeness of the bins. “+” in the right bubble plot indicates that the MAG could no longer be identified. “*” are MAGs where the species are known: Bin_68 (*Candidatus Syntrophosphaera thermopropionivorans*), Bin_67 (*Halopseudomonas salegens*), Bin_53 (*Candidatus Methanoculleus thermohydrogenitrophicus*), Bin_5 (*Acidilutibacter cellobiosedens*), Bin_25 (*Candidatus Brevifilum fermentans*) and Bin_22 (*Fermentimonas caenicola*) (also Supplementary Table 6).

Upon closer examination of the protein functions from the enriched MAGs, a notable abundance of stress proteins (up to 100% in some MAGs) was observed. Although ethanol can have a positive effect on methanogenesis in AD [55], the addition of ethanol in combination with the transfer of the MC from continuous LBR to serum bottles and mixing with a medium may have induced the expression of these stress proteins [56, 57]. Consequently, stress proteins were excluded from further analyses to ensure a metabolic enrichment of MAGs (Figure 5). In the case of the high-abundant MAGs, the protein abundance after the removal of stress proteins has only decreased slightly from Bin_41. However, in the case of the low-abundance MAGs, there have been some significant changes. MAGs such as Bin_74 (family *Desulfobacteraceae*), Bin_46 (unknown *Bacteria*), and Bin_20 (genus *Castellaniella*) showed more than 10-fold protein enrichment using the AA method. In contrast, proteins from MAGs such as Bin_13 (family: *Solibacteraceae*), Bin_34 (class *Betaproteobacteria*), Bin_25 (species *Candidatus Brevifilum fermentans*) and Bin_62 (unknown *Bacteria*) could no longer be identified after the stress-induced proteins were removed from the analysis. Therefore, the addition of ethanol resulted in increased expression of metabolic proteins in Bin_74, Bin_46, and Bin_20, whereas MAGs such as Bin_13, Bin_34, Bin_25, and Bin_62 primarily synthesized stress proteins.

### Ethanol-associated functions based on nP enrichment

Upon closer examination of Bin_46, Bin_74, and Bin_20, it was discovered that Bin_74 and Bin_20 expressed gluconeogenesis proteins. Too few proteins could be identified from Bin_46 to make conclusions about its metabolism. A review of the individual genes revealed that each of the three MAGs possesses at least two genes that encode for a potential alcohol dehydrogenase (ADH). Additionally, *Castellaniella* (Bin_20) is known to utilize acetyl-CoA from acetate, ethanol, or pyruvate *via* the glyoxylate cycle [58–62]. The incubation period for these three strains in the presence of ethanol was likely insufficient for the strains to multiply sufficiently to allow for the detection of these proteins. Without enrichment with the AA method, they would thus not have been the focus of analyses. In follow-up studies, the incubation times with ethanol could be extended so that also the metabolisms can be analysed [63].

The analysis of the Functional Ontology Assignments for Metagenomes in comparison to a sample with glucose (as in the LBR) showed that there were clear differences due to the substrate shift (Supplementary Figure 6). A clustered heatmap of the Functional Ontology Assignments for Metagenomes and most abundant MAGs showed that Bin_27, Bin_24, Bin_41, and Bin_53 in particular account for most of the functions of the microbial community of both replicates (Figure 6 and Supplementary Figure 7).

**Figure 6:**
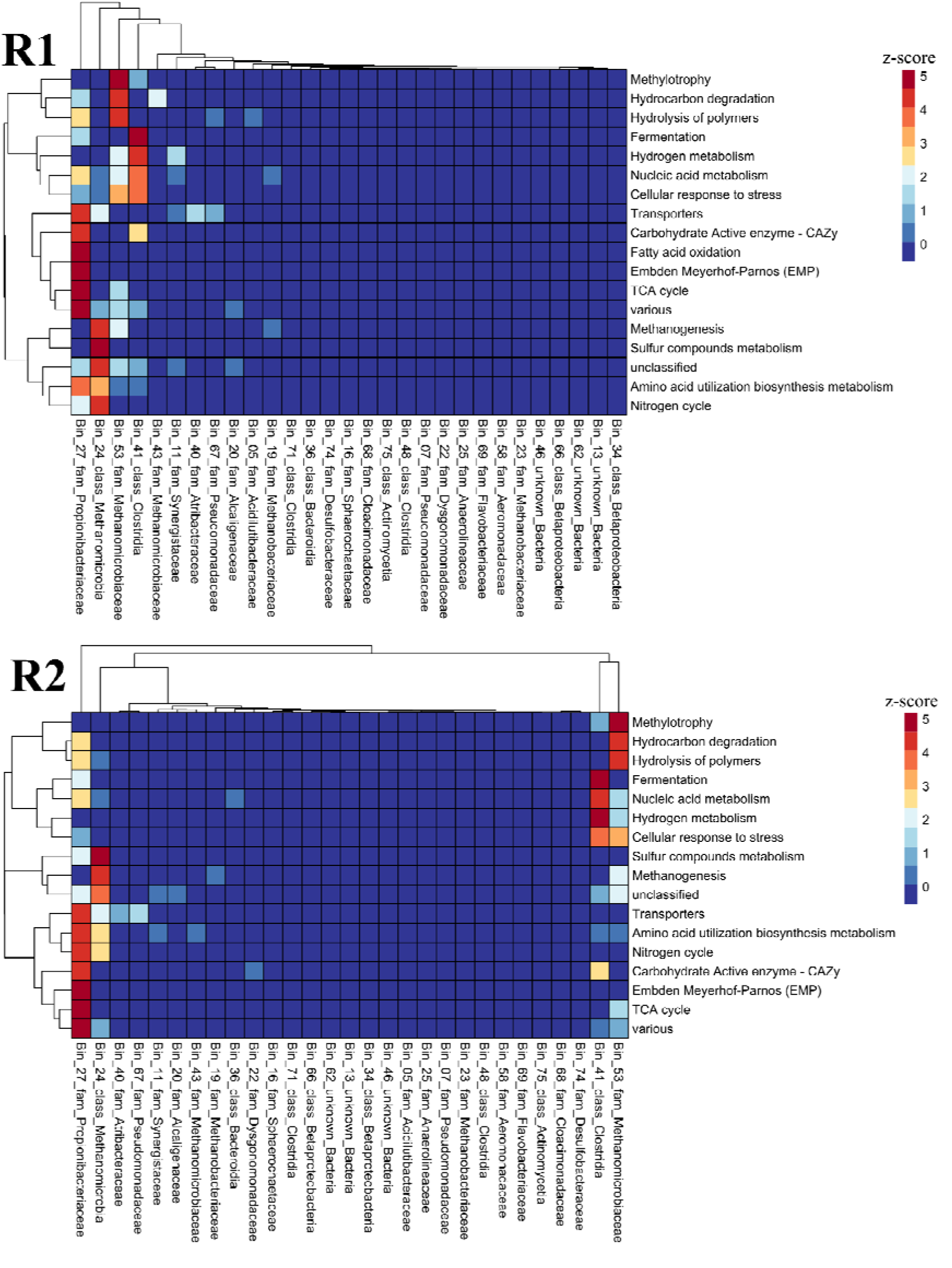
Alternated biological processes in the MC due to ethanol addition The functional ontology assignments for metagenomes of all enriched nP and MAGs with the AA method were analysed using a heatmap created with R Studio(2023.06.0 Build 421) and pheatmap (1.0.12). Previously, the protein abundances were normalised by dividing the spectral count of each protein by the total spectral abundance of the sample. Afterwards, a pivot table was generated, to sum up the relative abundances of the proteins based on the functional assignment and the MAG. The functional ontology assignments were normalized using a z- score per row (overall MAGs). also Supplementary Table 5.

A detailed analysis of proteomes of all samples identified 29 different ADHs, with 7 ADHs (Bin_41, 2x Bin_27, Bin_66, Bin_24, Bin_70, and Bin_53) as nP. Out of these 7 ADHs, 3 poorly classified ADHs (FHCFJDLK_121438 (Bin_41), FHCFJDLK_86872 (Bin_53), and FHCFJDLK_294919 (Bin_70, only in replicate 1)) were increased (≥ 2.5-fold) after enrichment (Supplementary Table 3 and 4). An analysis with Alphafold [64, 65] and KofamKOALA revealed that all three poorly classified ADHs are responsible for ethanol degradation (see Supplementary Table 4). However, Bin_70 was a very low abundant MAG (0.13%). This MAG belongs to the genus *Tepidiphilus*, which is known as a secondary fermenter and syntrophic acetate oxidizer [66, 67]. The abundance of this MAG is however insufficient to definitively ascertain its metabolic processes in this MC. Based on the proteins, Bin_41 appears to be an acetate producer. All proteins for acetate formation from ethanol were found in the proteome of Bin_41 [68], except an aldehyde dehydrogenase, which is necessary for the further degradation of acetaldehyde [69–71]. It is therefore assumed that the ADH of Bin_41 is bifunctional, converting ethanol into acetaldehyde (EC: 1.1.1.1) and then directly to acetyl-CoA (EC: 1.2.1.10, Supplementary Figure 8) [72–75]. Bifunctionality poses a challenge in the annotation of KO numbers, as normally only one KO number is assigned to one protein. Note, that it was only through enrichment of nP that we became aware of the protein and were able to consider the possibility of bifunctionality. Furthermore, NADH dehydrogenases, which are crucial for replenishing depleted NAD+ levels during anaerobic ethanol oxidation, were enriched from Bin_41 [76]. The removed electrons are probably transferred to a membrane-bound cytochrome-coupled formate dehydrogenase, which utilizes free CO_2_ and protons for formate formation at iron-containing 2Fe-2S clusters [77]. Additionally, ATP appeared to be generated through substrate chain phosphorylation [77, 78] (Supplementary Table 4). However, a major portion of stored energy is probably consumed for the reduction of formate due to coupled proton influx [79].

Bin_53 (species *Candidatus Methanoculleus thermohydrogenitrophicus*) is closely related to *Methanoculleus*, which is known to use ethanol and secondary alcohols as electron donors for methanogenesis [80]. Ethanol is converted into acetate by a reaction that is coupled with NADP+ reduction, which is then recovered by NADPH oxidoreductase (EC 1.5.1.40) [81, 82] delivering reduction equivalents to hydrogenotrophic methanogenesis (Supplementary Figure 9). Since ethanol is not completely oxidized to CO_2_, an alternative source of CO_2_ is required for hydrogenotrophic methanogenesis [83, 84]. A formate dehydrogenase (EC 1.17.1.9) was identified that converts formate into CO_2_, which can then be used for hydrogenotrophic methanogenesis and feeds released reduction equivalents into hydrogenotrophic methanogenesis. *Methanoculleus* is also known to use formate [84]. A syntropic interaction between Bin_53 and Bin_41 is plausible with Bin_41 delivering formate to Bin_53 [85–87]. Furthermore, Bin_53 also should be able to consume H_2_ which could result in the inhibition of Bin_41 when accumulating in the medium [88–90]. But syntrophic transfer of H_2_ probably did not play a major role in the LBR MC since no hydrogenase capable of releasing H_2_ was detected in Bin_41, (Figure 7). The phenomenon of syntrophic ethanol oxidation between fermenting bacteria and methanogenic archaea observed here has been repeatedly observed in anaerobic digesters [28, 91–93].

**Figure 7:**
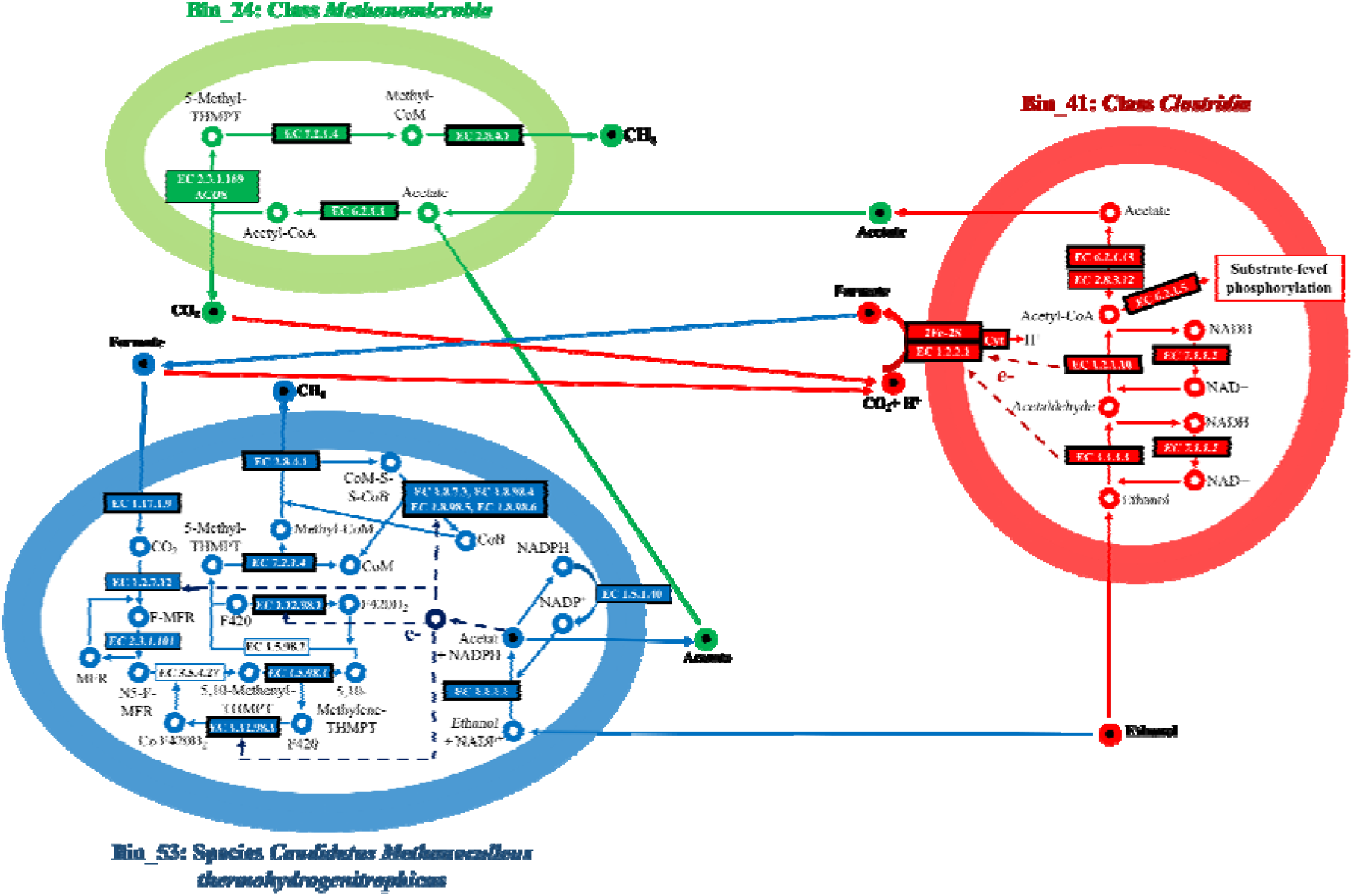
Suggestion for an anaerobic syntrophic ethanol oxidation based on nP enrichment. Intracellular metabolites are represented by unfilled circles, while extracellular metabolites are represented by filled circles. Metabolic pathways are designated by the colours of the arrows, which indicate the direction of metabolite reactions. Enzymes identified with the AA method are denoted by boxes on the arrows. The boxes with thicker lines indicate that the enzymes have been enriched at least twofold by the AA method. Exceptions include enzymes with EC numbers 1.5.98.2 and 3.5.4.27 that were not identified using the AA method or in the not enriched sample. The inspiration for this illustration is from Zhao et al. (2019) [100] Fig.7.

In Bin_24, all proteins necessary for acetoclastic methanogenesis were identified, indicating its potential to convert the acetate produced in Bin_41 and Bin_53 into methane and CO_2_ (Supplementary Figure 10 and Figure 7). Therefore, based on the enriched nP, a syntrophic relationship between these three MAGs is highly probable. In this scenario, ethanol is converted to acetate by Bin_53 and Bin_41, which transfer formate to each other, and Bin_24 consumes the acetate. However, other sources of H_2_, formate, acetate, and CO_2_, e.g. originating from fermentative degradation of microbial biomass [94], and additional partners involved in ethanol oxidation (e.g. by minor abundant bins) cannot be excluded (Supplementary Figures 11 and 12).

### Further Improvements and important considerations for implementing the workflow

A successful BONCAT experimentis the precondition for this workflow. Our experiments demonstrated the sufficient efficacy of BONCAT in slow-growing anaerobic MC. We advise reading Hatzenpichler et al [22, 23, 95] and Landor et al. [27] as they address pivotal aspects of a successful BONCAT experiment and the potential cytotoxicity of BONCAT. Following a successful BONCAT, our workflow can be readily implemented in any laboratory (SOPs in Supplementary Note 1). In the case of suboptimal nP yields despite a successful BONCAT experiment, increasing the bead volume is recommended. The sensitivity and reproducibility of quantitative measurements could be further improved by data-independent acquisition mass spectrometry measurements, such as DIA-PASEF^®^ [96]. In addition, fluorescence-activated cell sorting of ncAA-labelled cells from MC [24] could generate corresponding metagenomes for improved protein identification. Alternatively, the workflow could be adapted to HPG, pyrrolysine, or β-ethynylserine, although this has not yet been tested but the same biotin linker is also available for copper-based click chemistry [15, 97].

## Conclusion

The combination of BONCAT and the direct enrichment of nP is an effective approach for investigating MC using MS. Enriched nP reflects metabolic or adaptive processes, making it easier to get new insights into the MC by highlighting relevant proteins. The optimized nP enrichment workflow was demonstrated to be effective by a test system of *E. coli* and yeast and may serve as a reference for future nP enrichments. The application of the workflow on a slow-growing anaerobic MC enabled the description of syntrophic interaction in an MC consuming ethanol as substrate.

## Acknowledgment

We want to thank Dr.-Ing. Ute Thron from GETEC Green Energy GmbH for providing the fermentation substrate for the inoculum of the LBR.

## Supplementary

Supplementary note 1: SOPs and detailed method description

Supplementary note 2: additional figures and tables

Supplementary table 1: lab scale biogas reactor information

Supplementary table 2: protein data testsystem

Supplementary table 3: metaprotein data ethanol

Supplementary table 4: identified proteins for syntrophic interactions

Supplementary table 5: biolog processes

Supplementary table 6: input bubble plot

Supplementary table 7: raw data protein quantification

## Declarations

Institutional Review Board Statement: Not applicable.

Informed Consent Statement: Not applicable.

Data Availability Statement:

Proteome data were stored on PRIDE with the accession number: PXD047252, Username, Password: 0kWF7k9n

The metagenome database for protein identification is stored on ENA under the accession number: PRJEB70937

Conflicts of Interest: Not applicable.

Author contributions (CRediT):

- Conceptualization: P.H., D.B.
- Project administration: P.H., D.B.
- Formal analysis: P.H., D.K., D.W.
- Funding acquisition: U.R., R.H.
- Investigation: P.H., D.K., D.W.
- Methodology: P.H., D.K., D.W., T.B., A.W.
- Project administration: D.B., U.R.
- Software: -
- Resources: D.B., U.R., R.H.
- Supervision: D.B., U.R.
- Validation: PH., D.K., D.W., T.B., A.W.
- Visualization: P.H., D.K.
- writing—original draft: P.H.
- writing—review, and editing: D.K., R.H., A.D., D.W., U.R., D.B., T.B., A.W.

All authors have read and agreed to the published version of the manuscript.

## Abbreviations

AA: amino acids
AD: anaerobic digestion
ADH: alcohol dehydrogenase
AHA: 4-azido-l-homoalanine
BONCAT: bioorthogonal non-canonical amino acid tagging
BSA: bovine serum albumin
CC: click chemistry
DBCO: dibenzocyclooctyne
DTT: dithiothreitol
FASP: filter-aided sample preparation
HPG: homopropargylglycine
HPLC: high-pressure liquid chromatography
IAA: iodoacetamide
KEGG: Kyoto Encyclopedia of Genes and Genomes
LBR: laboratory biogas reactor
LC: liquid chromatography
MAG: metagenome assembled genome
MC: microbial community
MS: mass spectrometry
Mgf: mascot generic file
ncAA: non-canonical amino acids
nP: newly synthesized proteins
PASEF: parallel accumulation–serial fragmentation
PBS: phosphate-buffered saline
PEG: polyethylene glycol
rpm: rounds per minute
RT: room temperature
SDS: sodium dodecyl sulfate
SIP: stable isotope labeling
SOP: standard operation procedure
SS: disulfide
TYGS: Type (Strain) Genome Server

