## Supplementary Note 1 for "Tracing active members in microbial communities by BONCAT and click chemistry-based enrichment of newly synthesised proteins"

**Tracing active bugs in microbial communities by BONCAT and click chemistry-based enrichment of newly synthesised proteins**

*Patrick Hellwig^1,2*^, Daniel Kautzner^3^, Robert Heyer^3,4^, Anna Dittrich^5^, Daniel Wibberg^6,7^, Tobias Busche^8,9^, Anika Winkler^8,9^, Udo Reichl^1,2^, Dirk Benndorf^1,2,10*^*

*:	corresponding author

1. Otto-von-Guericke University Magdeburg, Bioprocess Engineering, Universitätsplatz 2, 39106 Magdeburg, Germany;
2. Bioprocess Engineering, Max Planck Institute for Dynamics of Complex Technical Systems, Sandtorstraße 1, 39106 Magdeburg, Germany
3. Multidimensional Omics Analyses group, Faculty of Technology, Bielefeld University, Universitätsstraße 25, 33615 Bielefeld,
4. Multidimensional Omics Analyses group, Leibniz-Institut für Analytische Wissenschaften – ISAS – e.V., Bunsen-Kirchhoff-Straße 11, 44139 Dortmund
5. Department of Systems Biology, Institute of Biology, Otto-von-Guericke University Magdeburg, , Universitätsplatz 2, 39106 Magdeburg, Germany
6. Institute for Genome Research and Systems Biology, CeBiTec, Bielefeld University, Universitätsstraße 25, 33615 Bielefeld, Germany
7. Institute of Bio- and Geosciences IBG-5, Computational Metagenomics, Forschungszentrum Jülich GmbH, 52425 Juelich, Germany
8. Center for Biotechnology - CeBiTec, Bielefeld University, Universitätsstraße 27, D-33615 Bielefeld, Germany.
9. Medical School East Westphalia-Lippe, Bielefeld University, Universitätsstraße 27, D-33615 Bielefeld, Germany.
10. Microbiology, Anhalt University of Applied Sciences, Bernburger Straße 55, 06354 Köthen, Germany

#### Table of contents:

SOP 1: BONCAT cell cultivation with *E. coli* and preparation of the test system	2

SOP 2: Click chemistry (in vitro)	7

SOP 3: Enrichment of newly synthesised proteins with magnetic streptavidin beads	12

SOP 4: Tryptic digestion using FASP (Filter Aided Sample Preparation)	19

SOP 5: SDS-PAGE	25

Full details of the other methods used	32

References	35

#### SOP 1: BONCAT cell cultivation with *E. coli* and preparation of the test system

#### Application:

An example of cell cultivation using BONCAT with AHA as a noncanonical amino acid and *E. coli* as the organism.

#### Comments:

- Do not use any reducing agents (like Dithiothreitol, 2-Mercaptoethanol, …) before executing click chemistry!

#### Abbreviations:

AHA	L-Azidohomoalanine

BONCAT	Bioorthogonal Noncanonical Amino Acid Tagging

RT	Room temperature

SDS	Dodecyl sulfate sodium salt

#### Chemicals:

Table 1: Chemicals used. Providing manufacturer and order numbers are not mandatory and can be replaced by other equivalent ones.

| **Chemicals** | **Manufacturer** | **Order Numbers** |
| --- | --- | --- |
| 4-Azido-L-homoalanine HCl (AHA) | Jena Bioscience | CLK-AA005-100 |
| Acetone | VWR | 20,066,330 |
| Ammonium chloride, NH_4_CL | VWR | 21236.291 |
| Calcium chloride * 2 H2O, CaCl | VWR | 22317.260 |
| Cyanase(TM) Nuclease solution | Serva Electrophoresis | 18542 |
| D(+)-Glucose | Sigma Aldrich | G-8270 |
| Di-Sodium hydrogene phosphate dihydrate, ≥99,5% p.a., Na_2_HPO_4_ | Roth | 4984 |
| Hydrochlorid acid 37%, HCl | VWR | 20,252,420 |
| Lactose | VWR | 101394S |
| Magnesium chloride (MgCl_2_) *6H_2_O | Sigma Aldrich | M9272-500g |
| Magnesium sulphate, pure, MgSO_4_ | Roth | 2611 |
| Milli-Q (ultrapure water) | / | / |
| Potassium chloride KCl, 99,5% | Roth | 6781.1 |
| Potassium dihydrogen phosphat, KH_2_PO_4_ | Merck | 1,048,731,000 |
| Silicia beads (0.5 mm) | BioSpec Products Inc | 1179105z |
| Sodium chloride, NaCl | Sigma | S9625 |
| Sodium dodecyl sulfate (SDS) | Serva | 20770.02 |
| Thiamine Hydrochloride (B1) | Sigma | T4625-25G |
| Tris | Applichem | A1086 |
| Yeast Extract | Merck | 1.11926.1000 |

#### Devices:

Table 2: Used devices. Devices and manufacturers are not mandatory and can be replaced by others with the same functionality.

| **Device** | **Name** | **Manufacturer** |
| --- | --- | --- |
| Ultrapure water extraction unit | Millipore Q-POD | Merck |
| FastPrep | FastPrep-96 | MP |
| Vortexer | Reax top | Heidolph |
| Centrifuge | Heraeus Multifuge 1S-R | ThermoScientific |
| Shaking incubator | 3033 | GFL |

#### Preparations and Solutions:

| **Solution** | **Chemicals** |
| --- | --- |
| **10 mM AHA solution** | 4.5 mg AHA  2.5 mL Milli-Q  sterile filtering  make fresh or set pH to 7 and store at 4°C for a short time |
| **20 mM Tris/HCL-MgCl buffer**  pH 7.5 | 0.24g Tris  80 mL Milli-Q  10 mg MgCl_2_  Adjust pH to 7.5 with HCl  Fill up to 100 mL with Milli-Q |
| **10% Lactose solution** | 10 g Lactose  100 mL Milli-Q |
| **10% Glucose solution** | 10 g Glucose  100 mL Milli-Q |
| **Thiamin solution** | 10 mg Thiamine  100 mL Milli-Q  Sterile filtering |
| **10% SDS** | 1 g SDS  Fill up to 10 mL with Milli-Q |
| **PBS**  1x PBS buffer | 8 g NaCl (137 mmol/L)  0.2 g KCl (2.7 mmol/L)  1.42 g Na_2_HPO_4_ (10 mmol/L)  0.27 g KH_2_PO_4_ (1.8 mmol/L)  1 L Milli-Q |
| **Incubation buffer I**  (PBS + 0.1 % SDS) | 99 mL PBS  1 mL 10% SDS |
| **0.1 M CaCl** | 3.8 g CaCl_2_  Add to 250 mL with Milli-Q |
| **1 M MgSO_4_** | 30,1 g MgSO4  Add to 250 mL with Milli-Q |
| **Ice-cold acetone** | 250 mL acetone  Store at -20°C |

| **Media** | **Chemicals** |
| --- | --- |
| **M9 media**  without carbon source | 6 g/L Na_2_HPO_4_  3 g/L KH_2_PO_4_  0.5 g/L NaCL  1 g/L NH_4_CL  **autoclave and then add:**  1 mL 1 M MgSO_4_  1 mL 0.1 M CaCl_2_ |
| **Yeast media** | 5 g/L Yeast extract  20 g/L Glucose |

#### Execution:

Organism used:

- *E. coli* (DSM 5911)
- Baker yeast (*S. cerevisiae*, DSM 1334)

#### 1 Cell cultivation and BONCAT

#### 1.1 *E. coli* pre-culture:

- Add 100 mL M9 medium (without carbon source) to a flask
  - Add 1 mL of 10% glucose solution (10 mL/L medium)
  - Add 100 µL thiamine solution (100mg/L medium)
- Inoculate flask with *E. coli*
- Incubate overnight at 37°C (Shaking incubator, 130 rpm)

#### 1.2 Yeast culture:

- Add 50 mL yeast medium to a flask
- Inoculate flask with *yeast*
- Incubate overnight at 30°C (Shaking incubator, 100 rpm)

#### 1.3 *E. coli* solution:

- Add *E. coli* pre-culture in a 50 mL reaction tube
  - Centrifuge at RT and 2,500 ×g for 10 min
  - Discard supernatant
  - Add the contents of the flask into the 50 mL reaction tube several times until the flask is empty
- Resuspend in 20 mL M9 medium (without carbon source)

#### 1.4 *E. coli* culture:

- Add 90 mL M9 medium to 2 flasks
  - Add 1 mL 10% lactose solution (10 mL/1 L medium)
  - Add 100 µL thiamine solution (100mg/1 L medium)
  - Add 1 mL of 10 mM AHA solution (100 µM AHA final concentration) to one flask (**AHA sample**) AND add 1 mL Milli-Q to another flask (**control**)
  - Add 10 mL *E. coli* solution in each flask (see 1.3)
  - Measure OD_600nm_ (should be approx. 0.1)
- Incubate at 37 °C for 2 h (Shaking incubator, 130 rpm)

#### 1.5 Yeast solution:

- Add the yeast culture in a 50 mL reaction tube
  - Centrifuge samples at 10,000 ×g, 4°C, 10 min
  - Discard supernatant
- Resuspend pellet in 20 mL 20 mM Tris/HCL-MgCl buffer (= **yeast solution**)

#### 1.6 Mixed *E. coli* and yeast sample:

- Harvest *E. coli* cultures (AHA sample and control)
  - Add flask content to a 50 mL reaction tube
  - Centrifuge at 10,000 ×g, 4°C, 10 min
  - Discard supernatant
  - Repeat until the flask is empty
- Add 10 mL yeast solution (see step 1.5) to each 50 mL reaction tube (with the pellets from the AHA sample and control)
  - Mix well
  - Centrifuge at 10,000 xg, 4°C, 10 min
  - Discard supernatant
- Store the pellets at -20°C or continue

#### 2. Cell disruption with a ball mill and lysis buffer

- Resuspend cell pellets in 5 ml 20 mM Tris/HCL-MgCl buffer
- Add 5 g silicia beads (1 g/mL)
- Add 2.5 μL Cyanase (TM) Nuclease solution (0.5 µl/mL)
- Shake at 1800 rpm for 5 min (FastPrep)
- Vortex and incubate at 37°C for 15 min
- Centrifuge at 8,500 ×g for 10 min at 4 °C
- Transfer supernatant to a new 50 mL reaction tube (proteins in solution!)

**3. Acetone precipitation:**

- Add the five-fold (v/v) amount of ice-cold acetone
- Incubate for at least 1 hour at -20°C (better overnight)
- Centrifuge at 8,500 ×g for 15-50 mL reaction tubes (16,400 ×g for 1-2 mL reaction tubes) for 30 min and 4°C
- Discard supernatant
- Dry the pellet (at RT under a hood)
- Resuspend pellet in **incubation buffer I**
- Optional: Incubate at 50°C for 1 h (If the pellet does not dissolve)
- Centrifuge for 10 min at 10,000 ×g
- Transfer supernatant to a new reaction tube

#### SOP 2: Click chemistry (in vitro)

#### Application:

Binding markers (biotin, fluorescent dye) to AHA-labelled proteins. The protocol is based on:

1. Hatzenpichler, R. & Orphan, V. J., 2015. Detection of Protein-Synthesizing Microorganisms in the Environment via Bioorthogonal Noncanonical Amino Acid Tagging (BONCAT). In: Hydrocarbon and Lipid Microbiology Protocols. Springer Protocols Handbooks. Berlin, Heidelberg: Springer Berlin Heidelberg, pp. 145-157. DOI: https://doi.org/10.1007/8623_2015_61

For this click chemistry method, high concentrations of strongly ionic reagents such as SDS (>0.2%) and high amounts of urea (>4 M) must not be used, as these have a negative impact on the reaction efficiency. See:

2. Dieterich, D. C., Lee, J. J., Link, A. J., Graumann, J., Tirrell, D. A., & Schuman, E. M. (2007). Labeling, detection and identification of newly synthesised proteomes with bioorthogonal non-canonical amino-acid tagging. Nat Protoc, 2(3), S. 532–540. DOI: 10.1038/nprot.2007.52
3. Yang, Y., Yang, X., & Verhelst, S. (2013). Comparative Analysis of Click Chemistry Mediated Activity-Based Protein Profiling in Cell Lysates. *Molecules, 18*(11), S. 12599-608. doi: 10.3390/molecules181012599

#### Comments:

- Do not use reducing agents before executing click chemistry!

#### Abbreviations:

AHA	L-Azidohomoalanine

BONCAT	Bioorthogonal Noncanonical Amino Acid Tagging

CY	Cyanin

IAA	2-Iodoacetamide

RT	Room temperature

SDS	Sodium dodecyl sulfate

DBCO	Dibenzocyclooctine

DMSO	Dimethyl sulfoxide

#### Chemicals:

Table 1: Chemicals used. Providing manufacturer and order numbers are not mandatory and can be replaced by other equivalent ones.

| **Chemicals** | **Manufacturer** | **Order Numbers** |
| --- | --- | --- |
| 2-Iodoacetamide (IAA) | AppliChem | A1666.0100 |
| 4-Azido-L-homoalanine HCl (AHA) | Jena Bioscience | CLK-AA005-100 |
| Acetone (≥99 %) | VWR | 20060321 |
| DBCO-Cy5.5 | Jena Bioscience | CLK-1046-1 |
| Dimethyl sulfoxide (DMSO) | Sigma-Aldrich | D8418 |
| Di-Sodium hydrogene phosphate dihydrate, ≥99,5% p.a., Na_2_HPO_4_ | Roth | 4984 |
| Disulfide Biotin DBCO (DBCO-SS-Biotin) | Click Chemistry Tools | A112-5 |
| Methanol (≥99,9 %) | Roth | T909.1 |
| Milli-Q (Ultrapure water) | / | / |
| Potassium chloride KCl, 99,5% | Roth | 6781.1 |
| Potassium dihydrogen phosphat, KH_2_PO_4_ | Merck | 1,048,731,000 |
| Sodium chloride, NaCl | Sigma | S9625 |
| Sodium dodecyl sulfate (SDS) | Serva | 20770.02 |

#### Devices:

Table 2: Used devices. Devices and manufacturers are not mandatory and can be replaced by others with the same functionality.

| **Device** | **Name** | **Manufacturer** |
| --- | --- | --- |
| Thermomixer | Thermomixer comfort | ThermoScientific |
| Ultrapure water extraction unit | Millipore Q-POD | Merck, Darmstadt, Germany |
| Centrifuge | Microstar 17R | VWR |

#### Preparations and Solutions:

| **Solution** | **Chemicals** |
| --- | --- |
| **PBS**  1x PBS buffer | 8 g NaCl (137 mmol/L)  0.2 g KCl (2.7 mmol/L)  1.42 g Na_2_HPO_4_ (10 mmol/L)  0.27 g KH_2_PO_4_ (1.8 mmol/L)  1 L Milli-Q |
| **10 % SDS solution** | 1 g SDS  Fill up to 10 ml with Milli-Q |
| **IAA solution**  200 mM Iodoacetamide (IAA) | 185 mg IAA  5 mL PBS  (prepare fresh before use) |
| **10 mM AHA solution** | 4.5 mg AHA  2.5 mL Milli-Q |
| **10 mM DBCO-CY5.5 stock solution**  Solved in Dimethyl sulfoxide (DMSO) | 1 mg DBCO CY5.5  85 µL DMSO |
| **250 nM DBCO-CY5.5 solution** | 1 μL 10 mM DBCO CY5.5 stock solution  1 mL DMSO |
| **10 mM DBCO-SS-Biotin stock solution** | 5 mg DBCO-SS-Biotin  575 µL DMSO |
| **3.75 µM DBCO-SS-Biotin solution** | 1.5 μL 10 mM DBCO-SS-Biotin stock solution  100 μL DMSO |
| **Incubation buffer I**  0.1 % SDS | 100 mL PBS  1 mL 10% SDS solution |

#### Execution:

It is recommended to do the click chemistry with fluorescent dye first to check the BONCAT efficiency (how many proteins are labelled with AHA). If the fluorescent signal is high, use 100 µg of protein for DBCO Biotin tagging; if the signal is low, use 200 µg of protein for DBCO Biotin tagging. If the signal is really low (with a small or no difference between the control and sample), the BONCAT labelling should be done again and optimized.

A few suggestions to increase BONCAT efficiency:

- Increase AHA concentration
- Lower incubation time when pulsed labelling is used
- Continuously add AHA when long incubation times are needed
- Check whether there is methionine in the incubation medium (methionine inhibits the AHA labelling)

#### 1.1: Tagging with a fluorescent dye (DBCO-CY5.5)

- Add 10 µg protein into a 2 mL reaction tube
- Fill up to 200 µL with incubation buffer I
- Add 200 µL IAA solution (final IAA concentration: 100 mM; final volume: 400 µL)
- Dark incubation for 1 h at RT and 300 rpm (Thermomixer)
- Add 10 µL 250 nM DBCO CY5.5 solution (dark environment)
- Dark incubation for 1 h at 37°C and 300 rpm (Thermomixer)
- End incubation by adding 40 µL 10 mM AHA solution
- Continue with **2. Acetone precipitation** **and methanol washing**
- Afterwards, do **SDS-PAGE** and **fluorescence scan** of the gel

#### 1.2: Tagging with biotin (DBCO-SS-Biotin)

- Add 100 - 200 µg protein into a 2 mL reaction tube (depending on BONCAT efficiency)
- Fill up to 200 µL with incubation buffer I
  - If sample volume > 200 µL: acetone precipitation with 110% of the protein amount and solubilize protein pellets in 200 µL incubation buffer I
- Add 200 µL IAA solution (final IAA concentration: 100 mM; final volume: 400 µL)
- Dark incubation for 1h at RT and 300 rpm (Thermomixer)
- Add 10 µL 3.75 µM Disulfide Biotin DBCO solution (dark environment)
- Dark incubation for 1h at 37°C and 300 rpm (Thermomixer)
- End incubation by adding 40 µL 10 mM AHA solution
- Continue with **2. Acetone precipitation and methanol washing**

#### 2: Acetone precipitation and methanol washing

- Add 1.5 mL ice-cold (-20 °C) acetone
- Incubation for at least 1 h at -20°C
- Centrifuge at 16,000 ×g for 30 min at 4°C
- Discard supernatant
- Wash 3 times with 1 mL methanol:
  - Centrifuge at 8,500 ×g for 10 min at 4°C or RT
  - Discard supernatant (it is not necessary to remove the entire liquid)
- Dry the pellet under the hood

#### SOP 3: Enrichment of newly synthesised proteins with magnetic streptavidin beads

#### Application:

Enrichment of biotin-marked AHA-labelled proteins with magnetic streptavidin beads. Some steps have been adapted from the following source:

1. Zhang, G. et al., 2014. In-Depth Quantitative Proteomic Analysis of de Novo Protein Synthesis Induced by Brain-Derived Neurotrophic Factor. Journal of Proteome Research, 13(12), pp. 5707-5714. DOI: <https://doi.org/10.1021/pr5006982>

#### Comments

- Carried out after cell cultivation with AHA (BONCAT) and tagging with Disulfide Biotin DBCO using click chemistry.
- Invitrogen™ Dynabeads™ MyOne™ Streptavidin C1 (and magnetic beads in general) can be destroyed or degraded by DTT (and in general by reducing agents). DTT can interact with the iron ions in magnetic beads, compromising their integrity and magnetic properties. Therefore, it should only be used in the elution step. During incubation with DTT, streptavidin can be released, and some beads may lose their magnetic properties, making it difficult to separate them from the sample. However, the FASP digestion, which should be used to digest the proteins afterward, filters out the beads. When maintained at 37°C, the number of beads that lost their magnetic properties was low.

#### Abbreviations:

AHA	L-Azidohomoalanine

BONCAT	Bioorthogonal Noncanonical Amino Acid Tagging

BSA	Albumin, from bovine serum

DTT	Dithiothreitol

RT	Room temperature

SDS	Sodium dodecyl sulfate

#### Chemicals

Table 1: Chemicals used. Providing manufacturer and order numbers are not mandatory and can be replaced by other equivalent ones.

| **Chemicals** | **Manufacturer** | **Order number** |
| --- | --- | --- |
| 1,4-Dithiothreitol (DTT) | Roth | 6908.2 |
| Acetone (≥99 %) | VWR | 20060321 |
| Acetonitrile (MS/MS grade) | VWR | #83640.320 |
| Albumin, from bovine serum (BSA) | Sigma | A3912 |
| Glutamic acid | Merck | 2.910.250 |
| Glutamine | Sigma Aldrich | G3126 |
| Glycerol | Carl Roth | 3783.1 |
| Glycerol | Roth | 3783.2 |
| Glycine | AppliChem | A3707,1000 |
| Histdidine | AppliChem | A1341,0100 |
| Hydrochlorid acid 37%, HCl | VWR | 20,252,420 |
| Invitrogen™Dynabeads™ MyOne™ Streptavidin C1 | Invitrogen™ | 65001 |
| LC-MS grade water | VWR | #83645.320 |
| Leucine | AppliChem | A1426 |
| Mercaptoethanol (toxic ☠☠☠) | Sigma | M3148 |
| Milli-Q (Ultrapure water) | / | / |
| Potassium chloride KCl, 99,5% | Roth | 6781.1 |
| Potassium dihydrogen phosphate, KH_2_PO_4_ | Merck | 1,048,731,000 |
| Sodium chloride, NaCl | Sigma | S9625 |
| Sodium dodecyl sulfate (SDS) | Serva | 20770.02 |
| Sodium hydroxide, NaOH | VWR | 28248298 |
| Tris | Applichem | A1086 |
| Tryptophan | Merck | 1.083.740.100 |
| Tween 20 | Roth | 9127.1 |
| Urea | AppliChem | A1049.1000 |

#### Devices:

Table 2: Used devices. Devices and manufacturers are not mandatory and can be replaced by others with the same functionality.

| **Device** | **Name** | **Manufacturer** |
| --- | --- | --- |
| Thermomixer | Thermomixer comfort | ThermoScientific |
| Magnetic rack | MagRack6 | Jena Bioscience |

#### Preparations and Solutions:

Table 3: List of Solutions and required chemicals.

| **Solution** | **Chemicals** |
| --- | --- |
| **PBS**  1x PBS buffer pH 7.4 | 8 g NaCl (137 mmol/L)  0.2 g KCl (2.7 mmol/L)  1.42 g Na_2_HPO_4_ (10 mmol/L)  0.27 g KH_2_PO_4_ (1.8 mmol/L)  1 L Milli-Q |
| **10% SDS solution**  0.1 g/ml SDS | 1 g SDS  Fill up to 10 ml with Milli-Q |
| **5 M NaOH** | 20 g NaOH  100 mL Milli-Q |
| **0.1 M Tris-HCl**  pH 8.5 | 15.7 g Tris-HCl  800 mL Milli-Q  Adjust pH value to 8.5 (add 5 M NaOH)  Fill up to 1 L with Milli-Q |
| **Incubation buffer III**  PBS + 0.1% SDS + 0.1% Tween 20 | 50 mL PBS  500 µL 10% SDS-solution  50 µL Tween 20  (prepare before use) |
| **BSA block solution** | 10 mg BSA  10 mL Incubation buffer III |
| **Amino acid block solution** | 0.5 mL 1 mg/mL Leucine in PBS  0.5 mL 1 mg/mL Histidine in PBS  0.5 mL 1 mg/mL Tryptophan in PBS  0.5 mL 1 mg/mL Glycine in PBS  0.5 mL 1 mg/mL Glutamine in PBS  0.5 mL 1 mg/mL Glutamic acid in PBS  27 mL Incubation buffer III  (prepare before use) |
| **Washing buffer I**  PBS + 0.1% SDS + 0.1% Tween 20 + 100 mM NaCl | 100 mL PBS  1 mL 10% SDS solution  100 µL Tween 20  0.5844 g NaCl  (prepare before use) |
| **Washing buffer II**  7 M Urea buffer | 63.063 g Urea  fill up to 150 ml with 0.1 M Tris-HCl (pH 8.5) |
| **Washing buffer III**  20% Acetonitrile | 16 mL LC-MS grade water  4 mL Acetonitrile  (prepare before use) |
| **Elution buffer**  50 mM DTT | 10 mL Incubation buffer III  77 mg DTT  (prepare before use) |
| **2% SDS-Puffer** | 80 mL PBS  20 mL 10% SDS solution |
| **0.5 M Tris-HCl**  pH 6.8 | 100 mL dest. water  15 g Tris  Adjust pH 6.8 with 1 M HCl  Fill to 250 mL with dest. water  Note the final pH value on the bottle |
| **SDS-sample buffer** | 51 mL Milli-Q  12.5 mL 0.5 M Tris-HCl, pH 6.8  10 mL glycerol  20 mL 10% SDS solution  5 mL mercaptoethanol (toxic ☠☠☠, fume hood) |

Table 4: Amino acids used for blocking and their property.

| **Amino acid** | **Side chain property** |
| --- | --- |
| Leucine | aliphatic hydrophobic |
| Tryptophan | aromatic hydrophobic |
| Histidine | polar positive |
| Glutamine | polar neutral |
| Glutamic acid | polar negative |
| Glycine | nonpolar neutral |

#### Execution:

For blocking with BSA, do step 3.1. For blocking with amino acids, do step 3.2, or neither if you are not blocking. **Blocking with amino acids works best and is recommended.**

#### 1. Resuspend proteins

- Resuspend proteins (see SOP click chemistry) in 400 µL **incubation buffer III**
  - 50°C for 30 - 60 min (Thermomix)
- Centrifuge at 16,400 ×g for 5 min

**In all following steps (until elution) ensure that the beads do not clump**

- **Carefully pipette up and down during each step**
- **If they clump: wash them with incubation buffer III**

#### 2. Preparing beads

- Vortex beads (Invitrogen™Dynabeads™ MyOne™ Streptavidin C1) for 30 sec
- Add 50 µL in a protein LoBind reaction tube
- Wash 2x with 500 µL LC-MS grade water

#### 3. Blocking the beads

Choose either Step 3.1 or 3.2 (recommended), or skip this step if you do not wish to block the beads.

##### **3.1 Blocking with BSA (not recommended)**

- Add 1 mL BSA block solution to the beads
- Incubate for 30 min at RT and 400 rpm (Thermomix)
  - Place the reaction tube on a magnetic rack (2-3 min) and discard the supernatant
- Wash 1x with 1 mL incubation buffer III
  - Place the reaction tube on a magnetic rack (2-3 min) and discard the supernatant

##### **3.2 Blocking with amino acids (recommended)**

- Add 1 mL amino acid block solution to the beads
- Incubate for 30 min at RT and 400 rpm (Thermomix)
  - Place the reaction tube on a magnetic rack (2-3 min) and discard the supernatant
- Wash 1x with 1 mL incubation buffer III
  - Place the reaction tube on a magnetic rack (2-3 min) and discard the supernatant

#### 4. Binding labelled proteins to beads

- Add sample (step 1) to the beads
  - Pipette up and down
- Incubation for 30 min at RT and 400 rpm (Thermomix)
  - Place the reaction tube on a magnetic rack (2-3 min) and discard the supernatant

#### 5. Washing

- Wash **3x** with 1 mL **washing buffer I**
  - Incubate for 1 min
  - Place the reaction tube on a magnetic rack (2-3 min) and discard the supernatant (or collect the supernatant for control later)
- Wash **3x** with 1 mL **washing buffer II**
  - Incubate for 1 min
  - Place the reaction tube on a magnetic rack (2-3 min) and discard the supernatant (or collect the supernatant for control later)
- Wash **3x** with 1 mL **washing buffer III**
  - Incubate for 3 min
  - Place the reaction tube on a magnetic rack (2-3 min) and discard the supernatant (or collect the supernatant for control later)
- Wash **2x** with 1 mL **washing buffer I**
- After washing with acetonitrile, the beads can be very clumpy
  - Incubation for 1 min
  - Place the reaction tube on a magnetic rack (2-3 min) and discard the supernatant (or collect the supernatant for control later)

#### 6. Eluting proteins

- Add 200 µL **elution buffer**
  - Contains **DTT (see comment!)**
- Dark incubate at 30 min at 37°C and 500rpm (Thermomix)
- Place reaction tube on magnetic rack (2-3 min)
  - **Supernatant** = **Enriched proteins** (Elution (E))
- Remove supernatant:
  - Break point: Add to a new protein lobind tube and at -20°C

OR

- Continue: Add onto a Centrifugal Filter Unit (10 kDa) and continue with tryptic digestion (SOP FASP digestion)

#### 7. Extra: test the elution quality – boiling beads

- Add 200 µL **2 % SDS-sample buffer**
  - Contains **reducing agent** mercaptoethanol **(see comment!)**
- Boil at 60 °C for 5 minutes
- Place reaction tube on magnetic rack (2-3 min)
- Add supernatant in a Centrifugal Filter Unit (10 kDa)
  - Supernatant = Sample: **Beads Boiled (BB)**

**Or:**

- Add 200 µL **2% SDS-Puffer**
- Boil at 96 °C for 5 minutes
- Place reaction tube on magnetic rack (2-3 min)
- Add supernatant in a Centrifugal Filter Unit (10 kDa)
  - Supernatant = Sample: **Beads Boiled (BB)**

#### SOP 4: Tryptic digestion using FASP (Filter Aided Sample Preparation)

#### Application:

Method for the tryptic digestion of proteins from protein extracts of biogas samples in liquid.

1. Wisniewski, J. R., Zougman, A., Nagaraj, N., and Mann, M. 2009. Universal sample preparation method for proteome analysis. Nature Methods, 6(5):359–362
2. Wiśniewski, J.R., Hein M.Y., Cox J., Mann M. Mol Cell Proteomics. 2014 Dec;13(12):3497-506. A "proteomic ruler" for protein copy number and concentration estimation without spike-in standards.

#### Abbreviations:

VE	Demineralised water

RT	Room temperature

BSA	Bovine Serum Albumin

M	Molarity

SDS	Sodium dodecyl sulfate

APS	Ammonium persulfate

LC	Liquid chromatography

MS	Mass spectrometer

TFA	Trifluoroacetic acid

DTT	1,4-Dithiothreitol

IAA	2-Iodoacetamide

#### Chemicals

Table 1: Chemicals used. Providing manufacturer and order numbers is not mandatory and can be replaced by other equivalent ones.

| **Chemicals** | **Manufacturer** | **Order number** |
| --- | --- | --- |
| 1,4-Dithiothreitol (DTT) | Roth | 6908.2 |
| 2-Iodoacetamide (IAA) | AppliChem | A1666.0100 |
| Acetic acid 99% (For LC-MS) | Acros | 295320025 |
| Acetonitrile (LC-MS grade) | VWR | 83640.320 |
| Ammonium hydrogen carbonate (ABC) | Carl Roth | T871.2 |
| Centrifugal Filter Units (10 kDa) | VWR | 516-8492 |
| Formic acid 99% (For LC-MS) | VWR | 20318.297 |
| Trifluoroacetic acid (TFA) (For LC-MS) | VWR | 84868.180 |
| Trypsin, Premium grade | Serva | 37286.04 (4x25μg) |
| Urea | AppliChem | A1049.1000 |
| Water (LC-MS grade) | VWR | 83645.320 |

#### Devices:

Table 2: Used devices. Devices and manufacturers are not mandatory and can be replaced by others with the same functionality.

| **Device** | **Name** | **Manufacturer** |
| --- | --- | --- |
| Thermomixer | *Thermomixer comfort* | ThermoScientific |
| Centrifuge | Microstar 17R | VWR |
| Vacuum centrifuge | *SPD1218* | ThermoScientific |

#### Preparations and Solutions:

Table 3: List of Solutions and required chemicals.

| **Solution** | **Chemicals** |
| --- | --- |
| **ABC buffer**  50 mM ABC, pH 7.8 | 98.80 mg ABC  25 mL Water (LC-MS grade)  (prepare before use) |
| **0.1 M Tris-HCl**  pH 8.5 | 15.7 g Tris-HCl  800 mL Water (LC-MS grade)  Adjust pH value to 8.5 (add 5 M NaOH)  Fill up to 1 L with Milli-Q |
| **8 M Urea buffer** | 4.8 g Urea  Fill up to 10 ml with 0.1 M Tris-HCl pH 8.5 |
| **400 mM DTT stock solution** | 24.68 mg DTT  400 μL 50 mM ABC buffer  (prepare before use and store in the dark) |
| **40 mM DTT working solution** | 0.4 mL 400 mM DTT stock solution  3.6 mL 8 M Urea buffer  (prepare before use and store in the dark) |
| **550 mM IAA stock solution** | 40.7 mg IAA  400 μL 50 mM ABC buffer  (prepare before use and store in the dark) |
| **55 mM IAA working solution** | 0.4 mL 550 mM IAA stock solution  3.6 mL 8 M Urea buffer  (prepare before use and store in the dark) |
| **Trypsin (25μg/25μl)** | Trypsin in 25 μl 50 mM acetic acid (c = 1 μg/μl) |
| **Extraction buffer**  5% Acetonitrile in ABC buffer | 1 ml 50 mM ABC buffer  50 µL Acetonitrile |
| **Solvent A (100% water + 0.1% TFA)** | 100 mL water (LC-MS grade)  0.1 mL TFA |

#### Execution:

Filters should be tested for functionality on the same day they are used. **Don't let them dry out (pay attention to this in every step).** Load the filter with 100 μl of Milli-Q, then centrifuge for 1 minute at 10,000 g. Check whether the liquid is evenly distributed at the top and bottom (if so, the filter is okay). If the liquid has flowed through completely, the filter is defective.

Samples that are already dissolved in the urea buffer should be adjusted to a volume of 200 μl with 8 M urea buffer. Then, transfer them to the filter (please skip to step 2.).

**1. Optional: Remove of SDS and Tween 20**

The FASP digestion should remove SDS and Tween 20 by washing steps with urea. To be on the safe side, SDS and Tween 20 can also be removed by acetone precipitation.

- Add five-fold (v/v) ice-cold acetone
- Incubation for at least 1 h at -20 °C
- Centrifuge at 16,000 ×g for 30 min at 4 °C
- Discard supernatant
- Dry the pellet under the hood
- Add 200 μL 8 M urea buffer
- Resuspend the protein

#### 2. Denaturation, reduction and alkylation

- Transfer protein solution to the filter unit (min vol. 50 μL / max vol. 500 μL)
  - Centrifuge at 10,000 ×g for 10 mins at RT (if there is still liquid on the filter, extend the centrifugation time)
- Wash with 200 μL 8 M urea buffer (Do it two times if you skipped step 1)
  - Centrifuge at 10,000 ×g for 10 min at RT (if there is still liquid on the filter, extend the centrifugation time)
- Add 100 μL of DTT working solution to the filter
  - Shake for 1 min at 800 rpm on the Thermomixer
  - Incubate at 56°C for 20 min with gentle shaking (300 rpm).
  - Centrifuge at 10,000 ×g for 10 mins at RT (if there is still liquid on the filter, extend the centrifugation time)
- Add 100 μL of IAA working solution to the filter
  - Shake for 1 min at 800 rpm on the Thermomixer
  - Incubate at 56°C for 20 min with gentle shaking in the dark (300 rpm).
  - Centrifuge at 10,000 ×g for 10 min at RT (if there is still liquid on the filter, extend the centrifugation time)
- Remove flow-through (save for later SDS-PAGE quality control or discard)
- Wash 1x with 100 μL urea buffer
  - Incubate for 2 min at RT on the Thermomixer
  - Centrifuge at 10,000 ×g for 5 min at RT (if there is still liquid on the filter, extend the centrifugation time)
- Wash 3x with 100 μL 50mM ABC buffer
  - Incubate for 2 min at RT on the Thermomixer
  - Centrifuge at 10,000 ×g for 5 min at RT (if there is still liquid on the filter, extend the centrifugation time)
- Discard flow-through

#### 3. Proteolytic digestion

- Prepare enzyme solution:
  - Dissolve 25 μL of lyophilized trypsin (Serva) in 25 μl of 50 mM acetic acid
  - Recommended enzyme:substrate ratio (E:S): 1:100 for trypsin
  - Prepare trypsin working solution in 50 mM ABC buffer, calculation: 200 μL per sample (For enrichment proteins 10 µg protein was assumed)
- Empty the lower part of the reaction vessel and wash with 250 µL ABC buffer (use a pipette)
- Add 200 µL enzyme solution to the filter
- Incubate for at least 2 h at 37°C with gentle shaking (300 rpm)

#### 3. Extraction

- Centrifuge at 10,000 ×g for 5 mins at RT **(DO NOT discard flow-through)**
- Add 50 µL extraction buffer to the filter
- Centrifuge at 10,000 ×g for 5 mins at RT **(DO NOT discard flow-through)**
- Add 50 µL Milli-Q (LC-MS grade) to the filter
- Centrifuge at 10,000 ×g for 5 mins at RT **(DO NOT discard flow-through)**
- Dispose of the filter unit (protein/peptide is in the flow-through!)
- Drying in a vacuum centrifuge
  - This should result in no liquid remaining, leaving only a peptide pellet
- Resuspend the pellet in solvent A (e.g. 1 µg/µL peptide concentration)
- Transfer to a MS-Vial

#### SOP 5: SDS-PAGE

#### Application:

SDS-PAGE enables protein separation for quality control. The protocol is according to:

1. Laemmli, U. K. (1970). "Cleavage of structural proteins during the assembly of the head of bacteriophage T4." Nature **227**(5259): 680-685.

#### Comments:

- Acrylamide and mercaptoethanol are toxic. Do all steps under a fume hood and wear glasses.
- For pH adjustment of Tris-HCl-solutions different HCl concentrations could be used. However, to low concentrations result in high volumes, which may exceed the maximum volume.
- Check whether you want to work with 1 mm or 1.5 mm thick gels and use corresponding glass plates, combs and gel volumes. The current during electrophoresis could be set below the maximum current. Also, duration of fixation or dyeing could be extended.
- The maximum/optimum loading volume per pocket for 1 mm gels is 30 μL/25 μL, 45 μL/40 μL for 1.5 mm gels, respectively.

#### Abbreviations:

APS	ammonium persulfate

dest. water	deionized water

DTT	1,4-Dithiothreitol

RT	room temperature

TEMED	tetramethylethyldiamin

SDS	sodium dodecyl sulfate

#### Chemicals

Table 1: Chemicals used. Providing manufacturer and order numbers are not mandatory and can be replaced by other equivalent ones.

| **Chemicals** | **Manufacturer** | **Order Numbers** |
| --- | --- | --- |
| Acetic Acid ≥ 96% | Carl Roth | T179.2 |
| Acetone, ≥ 99%., | VWR International | 20063.296 |
| Acrylamide solution 30% (toxic ☠☠☠) | Serva | 10688.01 |
| Ammonium sulfate | Carl Roth | 3746.3 |
| Ammonium persulfate (APS) | Merck | 1.01201.0500 |
| Bromophenol blue | GE Healthcare | 17-1329-01 |
| Coomassie BB G 250 | AppliChem | A3484.0100 |
| 1,4-Dithiothreitol (DTT) | Roth | 6908.2 |
| Ethanol (70%) | Carl Roth | T913.3 |
| Glycerol | Carl Roth | 3783.1 |
| Glycine | VWR International | 444495D |
| Hydrochloric acid (37%) | VWR International | 20252.420 |
| Isobutanol | AppliChem | A1150.1000 |
| Mercaptoethanol (toxic ☠☠☠) | Sigma | M3148 |
| Methanol ≥ 99.9% (toxic ☠☠☠) | Carl Roth | T909.1 |
| Protein standard | Thermo Scientific PageRuler Prestained | Protein Ladder #26616 |
| Phosphoric acid ≥ 85% | Carl Roth | 6366.2 |
| SDS (sodium dodecyl sulfate, (**toxic ☠☠☠**) | AppliChem | A2572.0250 |
| TEMED | Carl Roth | 2367.1 |
| Tris | AppliChem | A2264.1000 |

#### Devices:

Table 2: Used devices. Devices and manufacturers are not mandatory and can be replaced by others with the same functionality.

| **Device** | **Name** | **Manufacturer** |
| --- | --- | --- |
| Electrophoresis chamber | Mini-Protean Tetra System | BioRad (Hercules, USA) |
| Power supply | PowerPac Basic | BioRad (Hercules, USA) |
| Destilled water | Millipore Q-POD | Merck, Darmstadt, Germany |
| Centrifuge | Micro Star 17 R | VWR International, Darmstadt,  Germany |
| Centrifuge | Avanti J | Beckman Coulter, Brea, USA |
| Fluorescence scanner | *Odyssey Classic* | LI-COR |
| Scanner | *Biostep ViewPix 900* | EPSON |

#### Preparations and Solutions:

Table 3: List of Solutions and required chemicals.

| **Solution** | **Chemicals** |
| --- | --- |
| **5x SDS-running buffer** | 15 g tris  72 g glycine  5 g SDS  Fill to 1 L with dest. water  (Store at 4 °C) |
| **1x SDS-running buffer** | 200 mL 5x SDS-running buffer  800 mL dest. water |
| **SDS-solution**  0.1 g/mL SDS | 1 g SDS (toxic ☠☠☠)  Fill to 10 mL with dest. water  (Store at RT) |
| **1.5 M Tris-HCl**  pH 8.8 | 100 mL dest. water  90.75 g Tris  Adjust pH 8.8 with 4 M HCl  Fill to 500 mL with dest water  (Store at 4 °C) |
| **0.5 M Tris-HCl**  pH 6.8 | 100 mL dest. water  15 g Tris  Adjust pH 6.8 with 1 M HCl  Fill to 250 mL with dest. water  Note final pH value on the bottle  (store at 4 °C) |
| **Bromophenol blue-solution**  5 mg/ml bromophenol | 100 mL dest. water  0.5 g bromophenol blue  (Store at RT) |
| **SDS-sample buffer** | 50 mL dest. water  12.5 mL 0.5 M Tris-HCl, pH 6.8  10 mL glycerol  20 mL SDS-solution (0.1 g/mL)  5 mL mercaptoethanol (toxic ☠☠☠, fume hood)  1 mL bromophenol blue-solution (5 mg/ml)  (Store at 4 °C) |
| **APS-solution**  0.1 g/mL APS | 10 mL dest. water  1 g APS  Prepare aliquoits à 1 mL  (Store at -20 °C) |
| **Water-saturated butanol** | 100 mL dest. water  100 mL butanol  (Top phase is butanol) |
| **Fixation solution** | 329 mL dest. water  571 mL ethanol (70%)  100 mL acetic acid (100%) |
| **Coomassie Brilliant Blue Stock Solution** | 5 g Coomassie Brilliant Blue G-250  100 mL dest. water  (store at RT) |
| **Colloidal Coomassie Stock Solution** | 50 g ammonium sulfate  484 mL dest. water  6 mL ortho-phosphoric acid (85%)  10 mL Coomassie Brilliant Blue Stock solution  (Store at RT, acid cupboard) |
| **Colloidal Coomassie Dye Solution** | 200 mL Colloidal Coomassie Stock Solution  50 mL methanol  (Store at RT, solvent cupboard) |

#### Execution:

#### 1.1 Sample preparation by diluting:

- Dilute your sample in at least the same amount of SDS-sample buffer
- Shake 5 min at 60 °C and 1.400 rpm on the Thermomix
- Centrifuge at 16,400 xg and RT for 10 min
- **For 1 mm Gel:** Load 20 µL sample into each pocket
- **For 1.5 mm Gel:** Load 40 µL sample into each pocket

#### 1.2 Sample preparation by precipitation:

- Precipitate 110% of required sample amount
- Add same volume of dest. water to your sample
- Precipitate with fivefold of ice-cold acetone (100%)
- Incubate at least 1 h or overnight
- Centrifuge at 16,400 xg and RT for 10 min
- Discard supernatant and dry pellet
- **For 1 mm Gel**: Add 22 µL SDS-sample buffer to the dried protein pellet
- **For 1.5 mm Gel**: Add 42 µL SDS-sample buffer to the dried protein pellet
- Shake 5 min at 60 °C and 1,400 rpm on the Thermomixer
- Centrifuge at 16,400 xg and RT for 10 min
- **For 1 mm Gel**: Load 20 µL sample into each pocket
- **For 1.5 mm Gel**: Load 40 µL sample into each pocket

#### 2. Preparation of the gels and run electrophoresis

- Clean glass plates and spacer with ethanol and a fuzz free wipe
- Insert both into the clips. Proof whether the plates terminate flat on the ground
- Assemble gel casting device
- Prepare **12% separation gel**

Table 4: Steps for preparing the 12% separation gel

| 1 mm Separation Gel [mL] | 1 Gel | 2 Gels | 4 Gels | 6 Gels | 8 Gels |
| --- | --- | --- | --- | --- | --- |
| VE-Water | 2.23 | 4.46 | 8.92 | 13.38 | 17.84 |
| 1.5 M Tris-HCl pH 8.8 | 1.67 | 3.34 | 6.68 | 10.02 | 13.36 |
| SDS (10%) | 0.07 | 0.14 | 0.28 | 0.42 | 0.56 |
| Polyacrylamide | 2.67 | 5.34 | 10.68 | 16.02 | 21.36 |
| APS | 0.05 | 0.10 | 0.20 | 0.30 | 0.40 |
| TEMED | 0.01 | 0.01 | 0.02 | 0.03 | 0.04 |
| Total volume | **6.70** | **13.39** | **26.78** | **40.17** | **53.56** |
| 1.5 mm Separation Gel [mL] | **1 Gel** | **2 Gels** | **4 Gels** | **6 Gels** | **8 Gels** |
| VE-Water | 3.35 | 6.70 | 13.40 | 20.10 | 26.80 |
| 1.5 M Tris-HCl pH 8.8 | 2.5 | 5.00 | 10.00 | 15.00 | 20.00 |
| SDS (10%) | 0.1 | 0.20 | 0.40 | 0.60 | 0.80 |
| Polyacrylamide | 4 | 8.00 | 16.00 | 24.00 | 32.00 |
| APS | 0.075 | 0.15 | 0.30 | 0.45 | 0.60 |
| TEMED | 0.0075 | 0.02 | 0.03 | 0.05 | 0.06 |
| Total volume | **10.03** | **20.07** | **40.13** | **60.20** | **80.26** |

- Overlay liquid gel with water-saturated butanol and let it polymerize for 30 min
- Remove butanol and wash it with dest. water, remove the remaining water with a fuzzy-free wipe
- Prepare **4% stacking gel**

Table 5: Steps for preparing the 4% stacking gel

| 1 mm Stacking Gel [mL] | 1 Gel | 2 Gels | 4 Gels | 6 Gels | 8 Gels |
| --- | --- | --- | --- | --- | --- |
| VE-Water | 2.03 | 4.06 | 8.12 | 12.18 | 16.24 |
| 0.5 M Tris-HCl pH 6.8 | 0.83 | 1.66 | 3.32 | 4.98 | 6.64 |
| SDS (10%) | 0.03 | 0.06 | 0.12 | 0.18 | 0.24 |
| Polyacrylamide | 0.43 | 0.86 | 1.72 | 2.58 | 3.44 |
| Bromophenol blue | 0.013 | 0.03 | 0.05 | 0.08 | 0.10 |
| APS | 0.025 | 0.05 | 0.10 | 0.15 | 0.20 |
| TEMED | 0.005 | 0.01 | 0.02 | 0.03 | 0.04 |
| Total volume | **3.363** | **6.73** | **13.45** | **20.18** | **26.90** |
| 1.5 mm Stacking Gel [mL] | **1 Gel** | **2 Gels** | **4 Gels** | **6 Gels** | **8 Gels** |
| VE-Water | 3.05 | 6.10 | 12.20 | 18.30 | 24.40 |
| 0.5 M Tris-HCl pH 6.8 | 1.25 | 2.50 | 5.00 | 7.50 | 10.00 |
| SDS (10%) | 0.05 | 0.10 | 0.20 | 0.30 | 0.40 |
| Polyacrylamide | 0.65 | 1.30 | 2.60 | 3.90 | 5.20 |
| Bromophenol blue | 0.02 | 0.04 | 0.08 | 0.12 | 0.16 |
| APS | 0.0375 | 0.08 | 0.15 | 0.23 | 0.30 |
| TEMED | 0.0075 | 0.02 | 0.03 | 0.05 | 0.06 |
| Total volume | **5.065** | **10.13** | **20.26** | **30.39** | **40.52** |

- Add comb air bubble free on the stacking gel (Pay attention to splashing solution, wear glasses)
- Remove comb after 30 min polymerization time and wash pockets with dest. water
- Assemble each 2 gels with the spacer outside into the electrophoresis chamber.
- Fill the chamber with SDS-running buffer (1x)
- Remove air bubbles below the gel by pipetting
- Fill space into the chambers also with SDS-running buffer (1x)
- Fill 10-20 mL sample into each pocket (30 µL into 1.5 mm gels) and add 2 µL of the protein standard to one pocket (note the name of the standard)
- Close the chamber and connect it to the power supply
- Adjust for each gel 10 mA current until the migration front entered the separation gel and then 20 mA current. Stop short before the migration front has left the gel.
- After disconnecting the power supply remove the gels
- Continue with fixation

#### 3. Fixation of the gels:

- Place gels after electrophoresis into marked bowls
- Add fixation solution for at least 1 h on a shaker with 20 rpm.
- Wash twice with dest. Water

#### 4. Fluorescence scan:

When fluorescent-tagged proteins are used, perform a fluorescent scan before staining the gel. If you are using the fluorescence scanner “*Odyssey Classic*” (LI-COR), refer to the settings provided in Table 6. Place the gels on the scanner free of bubbles and dust.

Table 6: Fluorescence scanner “*Odyssey Classic*” settings for fluorescence scan of a gel.

| **Option** | **Setting** |
| --- | --- |
| Quality | Medium |
| Intensity | 0,5 |
| Channel | 700 |
| Resolution | 169 |
| Wavelength (Detection of DBCO-Cy5.5) | 700 nm |

#### 5. Coomassie Staining

- Add Colloidal Coomassie Dye Solution overnight at RT on a shaker with 20 rpm
- Wash with dest. Water

#### 6. Scan with Biostep ViewPix900 (N.1.01)

If you are using the scanner “*Biostep ViewPix 900*” (EPSON), refer to the settings provided in Table 7. Place the gels on the scanner free of bubbles and dust.

Table 7: Scanner “*Biostep ViewPix 900*” settings for scan of a gel.

| **Option** | **Setting** |
| --- | --- |
| Modus | Transmission |
| Color Intensity | 48 Bit color |
| Brightness | 0 |
| Contrast | 0 |
| Gamma | 1,2 |
| Red level, Green levek, blue level | 100, 100, 100 |

Protein extraction with methanol and chloroform (for impure samples)

The proteins of the MC of the LBR were extracted with four-fold volume (v/v) of methanol, one-fold volume (v/v) of chloroform, and three-fold volume (v/v) of pure water and centrifuged (RT, 10,000 ×g, 5 min). After centrifugation, the upper phase was carefully removed by pipetting and discarded without disturbing the middle phase. Next, four-fold (v/v) methanol was added. The suspension was vortexed and centrifuged (RT, 10,000 ×g, 10 min), and the supernatant was discarded. The protein pellets were dried under a fume hood and resuspended in incubation buffer II (PBS + 1% SDS). The samples were centrifuged (RT, 10,000 ×g, 10 min), and the supernatant was transferred to new reaction tubes.

AHA-labelling of nP from MC of LBR

50 mL culture suspension was taken from an MC of LBR (total volume of 1 L, temperature 40°C, 75 rpm agitation, pH value 7.4 – 7.5, feed 1 mL/h biogas medium (Medium optimized for anaerobic digestion process, see Supplementary table 2), and an average gas composition of 52% CH_4_ and 29% CO_2_) using a 50 mL syringe. The cultures were transferred into nitrogen-flushed 100 mL serum bottles (with a magnetic stirring bar) under an anaerobic workbench (Bugbox; Baker Ruskinn). The cultures were diluted 1:1 (v/v) with 50 mL biogas medium (without glucose). Afterwards, the cultures were incubated under the anaerobic workbench for one hour. After one hour, 1 mL ethanol solution (1:10 (v/v) with water) was added to each culture. For BONCAT, 1 mL 20 mM AHA was added (final concentration 200 µM), while the same volume of water was added to the controls without AHA. In parallel controls with 3.8 mM glucose instead of ethanol were prepared (without AHA). All serum bottles were sealed with an airtight lid with a 30 mL airtight syringe attached to the outlet. The cultures were then incubated for 24 h at 40°C with gentle stirring. The pH of the cultures was measured, and the cells were harvested through centrifugation (4°C, 10,000 ×g, 10 min). The supernatant was discarded, and the cell pellet was washed twice with 5 mL 20 mM Tris/HCL buffer (pH 7.5), followed by centrifugation (4°C, 10,000 ×g, 10 min), after which the supernatant was again discarded.

LC-MS/MS measurements

The peptides were analysed by LC-MS/MS using an UltiMate® 3000 nano splitless reversed-phase nanoHPLC (Thermo Fisher Scientific, Dreieich) coupled online to a timsTOF™ pro 1 mass spectrometer (Bruker Daltonik GmbH, Bremen). All chemicals were used in LC-MS grade. The peptides were trapped on a trap column (Dionex Acclaim, nano trap column, 100 μmi.d. x 2 cm, PepMap100 C18, 5 μm, 100 Å, nanoViper) with a flow rate of 15 μL/min loading A buffer (100% water, 0.1% TFA). Afterwards, the peptides were loaded on Dionex Acclaim PepMap C18 RSLC Nano-reversed phase column (2 μm particle size, 100 Å pore size, 75 μm inner diameter, and 500 mm length) for a chromatographic separation. The column temperature was 60°C, and the flow rate of the nanoflow with a binary solvent A/B gradient (solvent A: 100% ultrapure water, 0.1% formic acid; solvent B: 100% acetonitrile, 0.1% formic acid) was 400 nL/min. 1 µg of proteins from not enriched samples was injected. For the elutions and other fractions, 2 µL of the test system and 6 µL for the biogas proteins were injected. The separation started with 2% solvent B for 5 min, and the proportion of solvent B was linearly increased over a 120 min gradient to 35%. Afterwards, the separation column was washed with 95% solvent B for 10 min and re-equilibrated with 5% loading B for 20 min. At the same time, the trap column was washed with 95% loading B buffer (100% acetonitrile, 0.1% TFA) for 10 min and equilibrated with 100% loading A buffer (100% water, 0.1% TFA) at a flow rate of 45 µl/min for 20 min. The positive measurement mode of the MS and a data-dependent MS/MS method were used to measure the peptides. The precursor ions were selected with Trapped Ion Mobility Separation (TIMS) and the acquisition method Parallel Accumulation Serial Fragmentation (PASEF) from 0.6-1.6 V s cm^-2^. The precursors were trapped in a quadrupole with a resolution of 50.000 and a mass-to-charge ratio (m/z) ranging from 100-1.700. Subsequently, the fragment ions were produced from the ten most intense precursors (charge 1 - 5) of each spectrum (10 PASEF MS/MS scans per cycle) with collision-induced dissociation (CID). The resulting fragment ions were analysed in a TOF analyser. The analysis takes place with an active exclusion of the same precursor after 1 spectrum for 24 seconds or a release if intensity/previous intensity exceeds 4.

Protein identification using Mascot and the MetaProteomeAnalyzer (MPA)

The timsTOF™ 1 mass spectrometer MS/MS measurement raw data files were processed by the Compass Data Analysis software (version 5.1.0.177, Bruker Corporation, Bremen, Germany) and converted into Mascot Generic Files (mgf). The files were uploaded to our Mascot Server (version 2.6) for the peptide spectrum matching with the following parameters: enzyme trypsin, one missed cleavage, monoisotopic mass, Unimod 4 (cysteine) as fixed modification, Unimod 35 (methionine) and Unimod 896 (methionine), as variable modifications, ± 0.02 Da precursor and ± 0.02 Da MS/MS fragment tolerance, 0^13^C and +2/+3/+4 charged peptide ions. For the test system, a defined UniProtKB/SwissProt database (08/05/2022) containing *E. coli* K12 (taxonomy_id:83333) and yeast (taxonomy_id:4932) proteins were used. A metagenome from the LBR was used for the biogas samples (see Illumina library preparation, MiSeq sequencing, and metagenome assembly). The search results were exported as Mascot data files (dat) and uploaded to the MPA software version 3.0 [1] for generating metaproteins.

Further analysis of the metaprotein data

The metaproteins were exported from the MPA software as CSV and uploaded to Prophane (version 6.2.6) [2] for BLAST analysis of the functional and taxonomic annotation. Functional Ontology Assignments for Metagenomes (FOAM) were used for KO number and functional annotation in the Kyoto Encyclopedia of Genes and Genomes (KEGG) and diamond blastp for the taxonomic annotation. For all unknown KO numbers after FOAM annotation, KofamKOALA [3] was used to identify possible KO numbers (see Supplementary table 3).

Illumina library preparation, MiSeq sequencing, and metagenome assembly for protein database

A metagenome of the LBR’s MC was created to improve the identification of proteins from the MC. For three replicates from the LBR, whole metagenome shotgun PCR-free libraries were constructed from 5 μg of metagenomics DNA using the Illumina TruSeq® DNA PCR-free sample preparation kit (Illumina, Eindhoven, Netherlands) according to the manufacturer's protocol as described before [4]. The three libraries were quality controlled by analysis on an Agilent 2000 Bioanalyzer using the Agilent High Sensitivity DNA Kit (Agilent Technologies, Santa Clara, CA, USA) for fragment sizes of 500-1000 bp. Sequencing was performed on the MiSeq platform (Illumina; 2 × 300 bp paired-end sequencing, v3 chemistry). After sequencing, adapters, and low-quality reads were removed by an in-house software pipeline, as described earlier [5].

In the next step, the tool Megahit (v1.1.1) [6] was used to assemble the pooled sequencing data of all replicates using k-mer sizes of 21, 29, 39, 59, 79, 99, 119, and 141 (iterative assembly) and default settings. In addition, the gene prediction tool Prodigal v.2.6.0 [7] was applied in metagenome mode and normal mode to predict genes on assembled contigs larger than 1 kb. The translated amino acid sequences of the predicted genes were then used for further analysis. Metagenomic binning was done as previously described [8, 9] with small modifications. In brief, raw reads were aligned to the corresponding assembled metagenome contigs using Bowtie 2 (v2.4.1 [10]). The resulting SAM files were processed through SAMtools (v1.0 [11]). For binning, MetaBAT2 (v2.12.1 [12]) was applied with default settings. Completeness, contamination, and strain heterogeneity of the Metagenome assembled genome (MAGs) were estimated with BUSCO (v5.7.0 [13]), using the bacterial-specific single-copy marker genes database (odb10).
