## Supplementary Note 2 for "Tracing active members in microbial communities by BONCAT and click chemistry-based enrichment of newly synthesised proteins"

**Tracing active bugs in microbial communities by BONCAT and click chemistry-based enrichment of newly synthesised proteins**

*Patrick Hellwig^1,2*^, Daniel Kautzner^3^, Robert Heyer^3,4^, Anna Dittrich^5^, Daniel Wibberg^6,7^, Tobias Busche^8,9^, Anika Winkler^8,9^, Udo Reichl^1,2^, Dirk Benndorf^1,2,10*^*

*:	corresponding author

1. Otto-von-Guericke University Magdeburg, Bioprocess Engineering, Universitätsplatz 2, 39106 Magdeburg, Germany;
2. Bioprocess Engineering, Max Planck Institute for Dynamics of Complex Technical Systems, Sandtorstraße 1, 39106 Magdeburg, Germany
3. Multidimensional Omics Analyses group, Faculty of Technology, Bielefeld University, Universitätsstraße 25, 33615 Bielefeld,
4. Multidimensional Omics Analyses group, Leibniz-Institut für Analytische Wissenschaften – ISAS – e.V., Bunsen-Kirchhoff-Straße 11, 44139 Dortmund
5. Department of Systems Biology, Institute of Biology, Otto-von-Guericke University Magdeburg, , Universitätsplatz 2, 39106 Magdeburg, Germany
6. Institute for Genome Research and Systems Biology, CeBiTec, Bielefeld University, Universitätsstraße 25, 33615 Bielefeld, Germany
7. Institute of Bio- and Geosciences IBG-5, Computational Metagenomics, Forschungszentrum Jülich GmbH, 52425 Juelich, Germany
8. Center for Biotechnology - CeBiTec, Bielefeld University, Universitätsstraße 27, D-33615 Bielefeld, Germany.
9. Medical School East Westphalia-Lippe, Bielefeld University, Universitätsstraße 27, D-33615 Bielefeld, Germany.
10. Microbiology, Anhalt University of Applied Sciences, Bernburger Straße 55, 06354 Köthen, Germany

Supplementary Table S1: Optical density at 600 nm of *E. coli* and yeast measured with a photometer (Thermo Scientific Genesys 10S UV-VIS)

| **OD_600_ of the overnight cultures** | | |
| --- | --- | --- |
|  | *E. coli* | yeast |
| OD_600_: | 0.473 | 1.526 |
| **OD_600_ of *E. coli* at substrate shift** | | |
| time | AHA | No AHA |
| 0 h (shift to lactose): | 0.348 | 0.350 |
| 2 h (haverest): | 0.470 | 0.467 |

Optimization of the AHA-labelling of *E. coli*

The AHA labelling efficiency of the *E. coli* from the test system was improved for the later enrichment protocol development. Therefore, the optimal harvest time (0-8 h) of *E. coli* after substrate shift and AHA addition was analyzed with a flow cytometer after tagging ethanol-fixated *E. coli* cells with DBCO-Alexfluor 488 according to Hatzenpichler et al. (2016) [1]. *E. coli* cells showed the proportionally highest fluorescence in the flow cytometer after 2-4 h incubation with AHA (Supplementary Figure 1). Thus, *E. coli* was harvested for the subsequent experiments after 2 h. The AHA concentration did not need to be optimized because Hatzenpichler et al. (2014) [2] showed that an AHA concentration of 0.1 mM is optimal for *E. coli*.

Next, we used LC-MS/MS to investigate whether there were significant differences in the proteome of *E. coli* due to the addition of AHA. An analysis of the protein data with the Mann–Whitney U test showed no significant differences (see Supplementary Figure 2). 3 [3] and 2 [2] have also observed no significant influence of ncAA labelling on the *E. coli* proteome in the used concentration. Since labelling with AHA itself does not induce differences in the *E. coli* proteome, we used the AHA-labelled *E. coli* cells in combination with unlabelled yeast cells as a test system for protocol development.

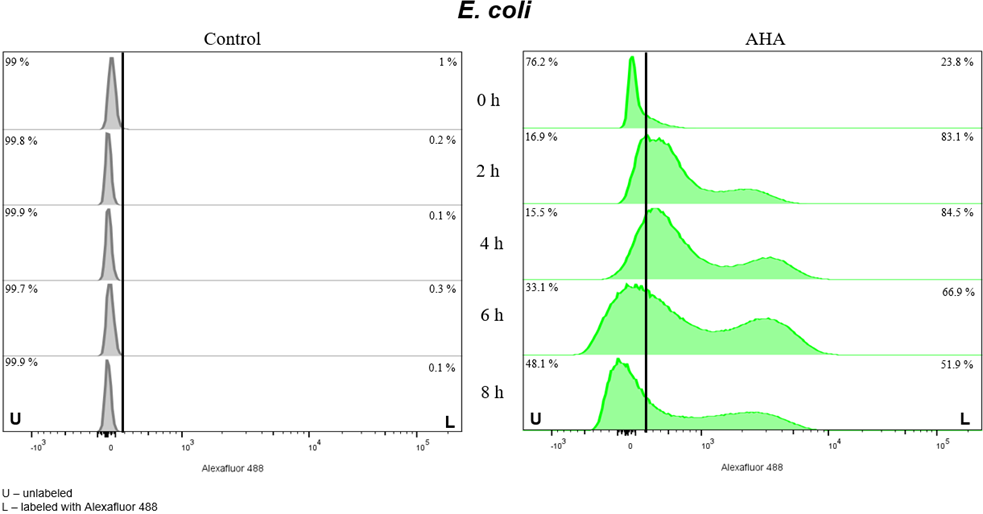

Supplementary Figure 1: Addition of a fluorophore to AHA-labelled *E. coli. E. coli* were incubated with AHA for the indicated times. *E. coli* was fixed with ethanol. Click-chemistry was used to label AHA-labelled *E. coli* with Alexafluor 488 (all steps according to [1]). Unlabelled /untreated *E. coli* served as control. *E. coli* was analyzed by flow cytometry using a FACS Canto II equipped with 3 lasers (405 nm, 488 nm, 663 nm), Firmware Version 1.47 (BD Biosciences, Franklin Lakes, NJ, USA). The threshold for background fluorescence of *E. coli* incubated without AHA at the starting point of the experiment was set to 1 % (vertical line in the histograms). Percentages of unlabelled (u) and Alexa-Fluor488-labelled *E. coli* (l) for each analyzed time-point are depicted.

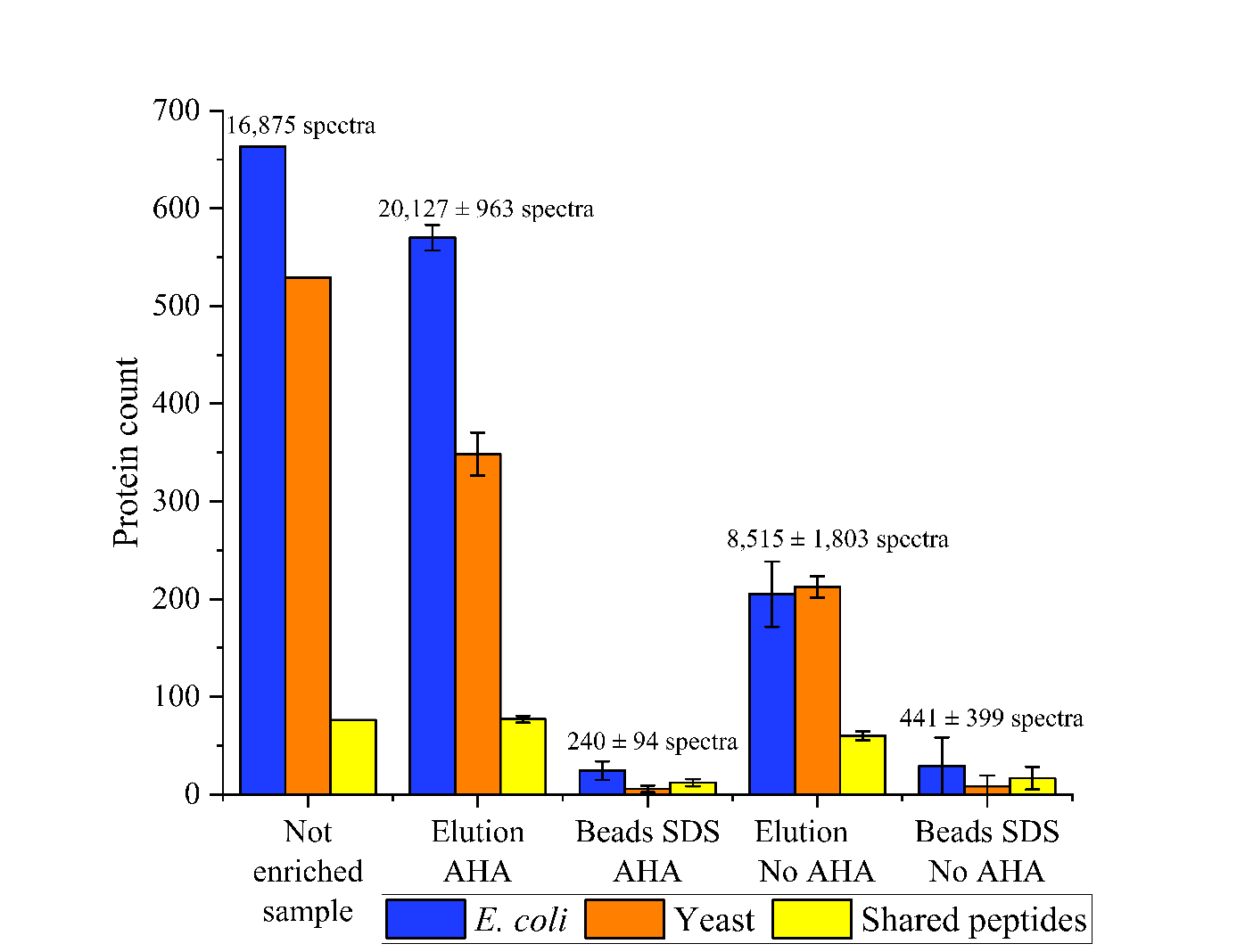

Supplementary Figure 2: Unsuccessful enrichment of newly synthesized proteins with the adapted protocol of the beads manufacturer and Zhang et al. (2014) [4]. The samples were handled as mentioned in the material and methods section. After binding the proteins to the beads, the beads were washed three times with PBS + 0.1% Tween20 + 0.1% SDS, three times with 7 M Urea + 0.1 M Tris-HCl pH 8.5, three times with 20% acetonitrile + 80% pure water and two times with PBS + 0.1% Tween20 + 0.1% SDS. Afterwards, the protocol mentioned in the material and methods section was used. The protocol resulted in a high background of yeast proteins (orange bar). Therefore, the protocol was optimized in this work. The bars represent the average of three technical replicates and the error bars represent the standard derivation.

**Comparison of the three tested enrichment methods**

Three strategies were examined, including a no blocking (nB) with high NaCl concentration (237 mM final concentration), blocking the beads with BSA (plus 237 mM NaCl, BSA-B), and blocking the beads with an amino acid mixture (plus 237 mM NaCl, AS-B) (see Supplementary Figure 4). The comparison aimed to select the best approach for stimulating nP. The proteins of the best approach should then be further investigated. The comparison of the approaches was based on three criteria: reliability, effectiveness, and specificity. The comparison was made for the elutions since only specific bound (AHA-marked) proteins should be enriched. The proteins assigned to “beads boiled” (see Supplementary Figure 5) are not considered in the comparison because they cannot be identified as *E. coli* or yeast proteins.

The identified proteins evaluate the reliability of the method. A (relative) standard deviation was used to determine the scatter between technical replicates. A total of 533 (nB), 365 (BSA-B), and 607 (AS-B) proteins were identified in the elution with AHA for the approaches. The standard deviation for the approaches was 17% (nB), 34% (BSA-B), and 3% (AS-B). Thus, the standard deviation of the AS-B Elution with AHA was about 6 times smaller than that of the nB and about 11 times smaller than that of the BSA-B. Thus, the AS-B approach had the highest reliability (see Supplementary Figure 4).

The effectiveness of the method was evaluated using the enriched *E. coli* proteins. For this purpose, the proteins in all replicates (at least 2 spectra per protein) were. Subtraction of the control proteins (Elution no AHA) yields the enriched *E. coli* proteins. After that, the AS-B approach enriched with 472, the most *E. coli* proteins. This was 84 more *E. coli* proteins than the nB approach and 235 more E. coli proteins than the BSA-B approach. Thus, the AS-B approach had the highest effectiveness (see Supplementary Figure 4).

The specificity of the method is evaluated by the number of yeast proteins and their spectra in the elution with AHA and the number of proteins in the control (elution no AHA). In the approaches, 12 (nB), 7 (BSA-B), and 21 (AS-B) yeast proteins were identified in the elution with AHA. For a total of 7,855 (nB), 4,452 (BSA-B), and 9,128 (AS-B) identified spectra, 0.50%, 0.45%, and 0.60% of the spectra were proportionally assigned to yeast (see Supplementary Figure 4). Due to the small number of spectra, the yeast proteins are negligible. In the controls (elution no AHA), a total of 26 (nB), 11 (BSA-B), and 10 (AS-B) proteins were identified. Of these, 8 (nB), 6 (BSA-B), and 3 (AS-B) proteins occur in all replicates. Overall, yeast proteins were negligible in all approaches.

Finally, it was checked whether the specific elution of the proteins worked completely. For this purpose, the boiled beads were examined. High temperatures lead to a denaturation of streptavidin, which releases the bound proteins [5]. Thus, after boiling, all proteins not eluted by DTT should now separate from the beads. The number of SDS eluted proteins (and spectra) for all approaches was the same as for the controls. Accordingly, adding DTT eluted all proteins specifically bound to the beads.

Blocking with amino acids (approach AS-B) identified significantly more proteins with a lower standard deviation compared to blocking with BSA (approach BSA-B) and the unblocked beads (approach nB). Therefore, blocking with amino acids appears to reduce steric effects significantly. In addition, blocking with amino acids also resulted in fewer proteins in the control, implying that fewer proteins bind nonspecifically to the mSB. Blocking with BSA (approach BSA-B) reduced nonspecific and specific binding proteins compared to unblocked beads (approach nB).

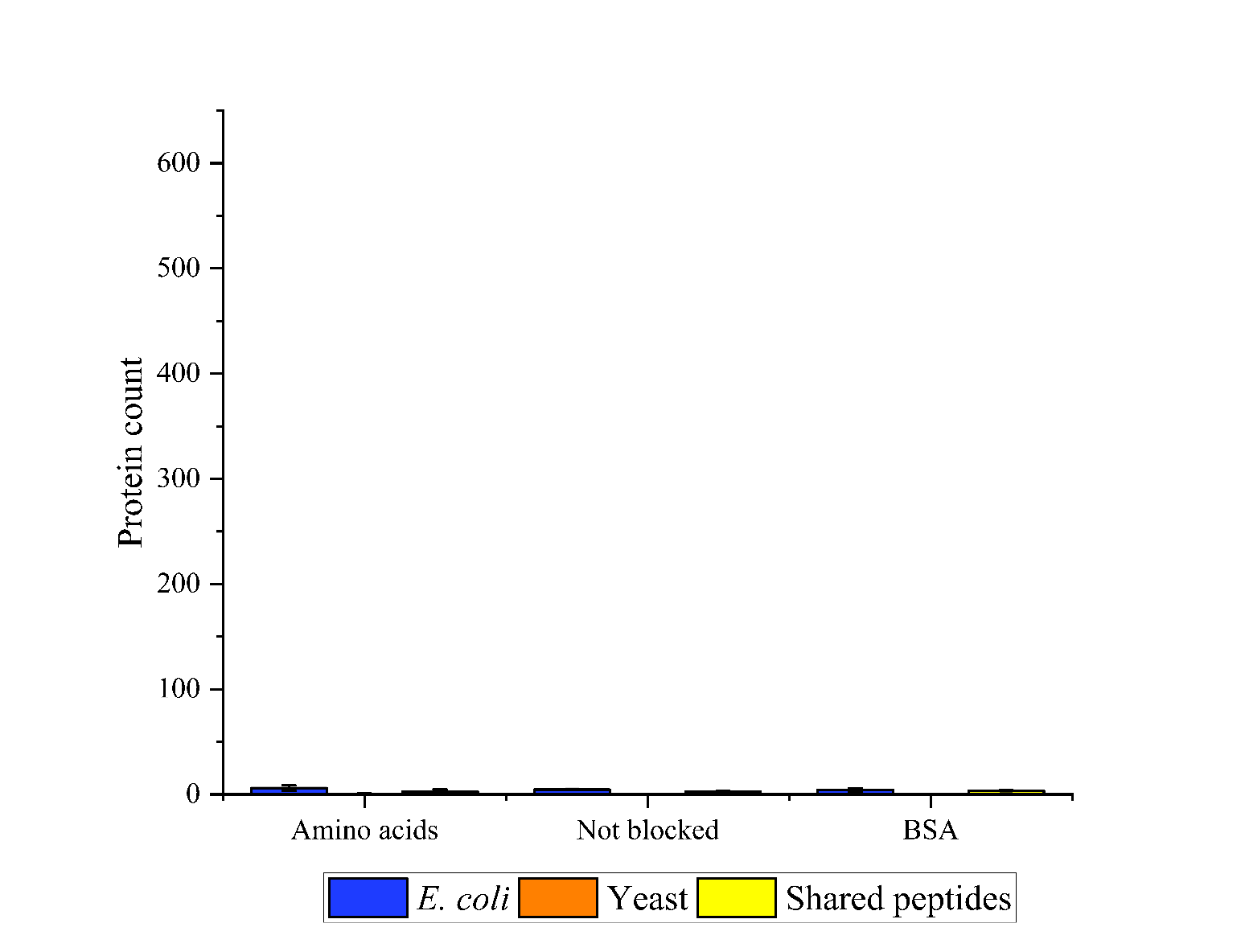

Supplementary Figure 3: Identified remaining proteins on the beads of the three enrichment methods after elution with DTT. After the elution of the nP with DTT, the beads were destroyed by boiling them in an SDS buffer. All remaining proteins on the beads were analyzed. All proteins with at least two spectra were considered. The proteins were grouped into the taxonomies *E. coli* and yeast. Proteins with shared peptides were grouped into “shared peptides”. The bars represent the average of each group and the error bars represent the standard derivation of three technical replicates.

Supplementary Table S2: Measured pH values of the biogas microbiome after 24 h incubation

| **Sample** | **pH** |
| --- | --- |
| AHA/- R1 | 7.5 |
| AHA/- R2 | 7.4 |
| AHA/- R3 | 7.4 |
| AHA/+ R1 | 7.3 |
| AHA/+ R2 | 7.4 |
| AHA/+ R3 | 7.4 |

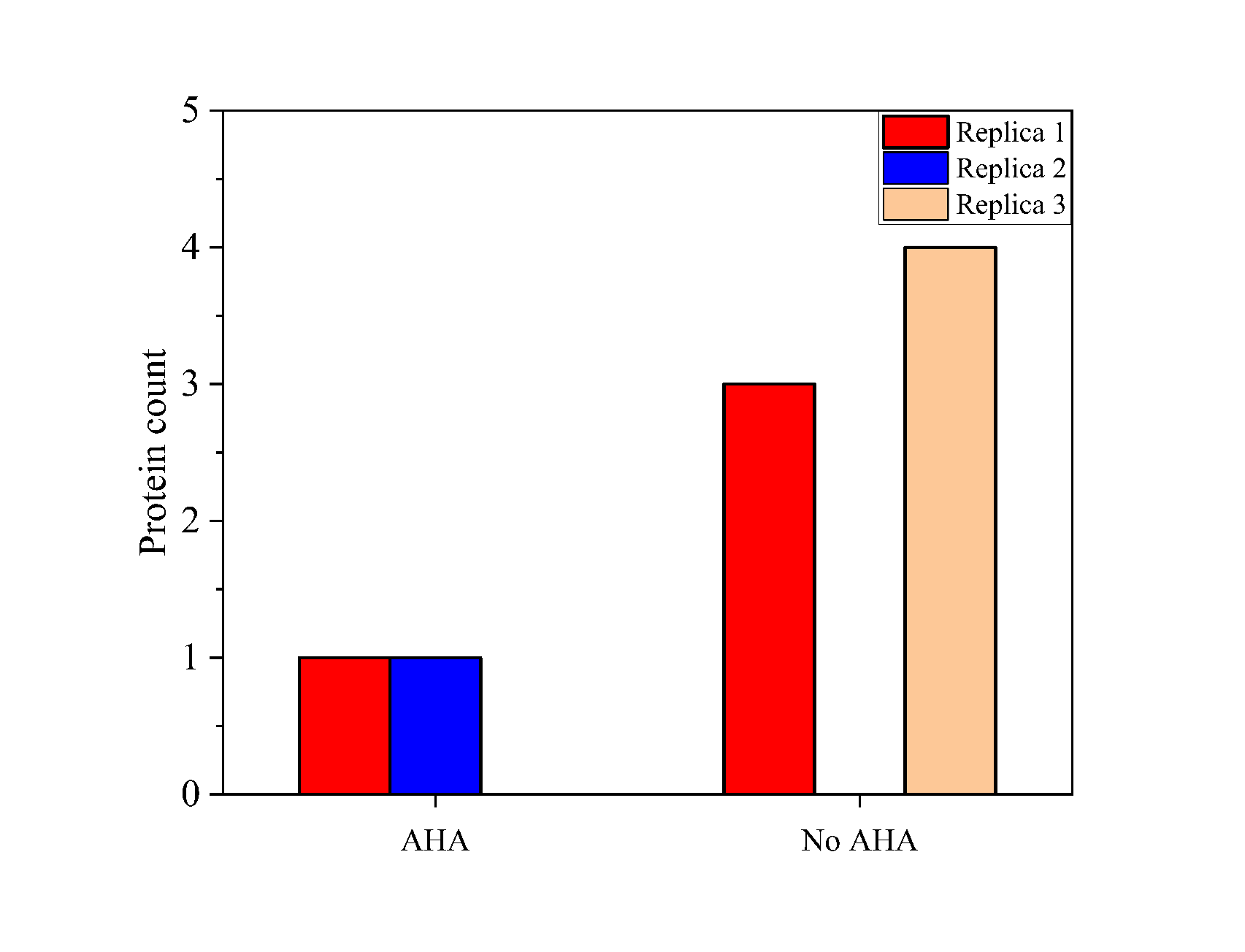

Supplementary Figure 4: Identified remaining proteins on the beads of nP enrichment from the samples of the microbial communities from the laboratory biogas reactor. After eluting the nP with DTT, the beads were destroyed by boiling them in an SDS buffer. All remaining proteins on the beads were analyzed. The bars represent the average of each group and the error bars represent the standard derivation.

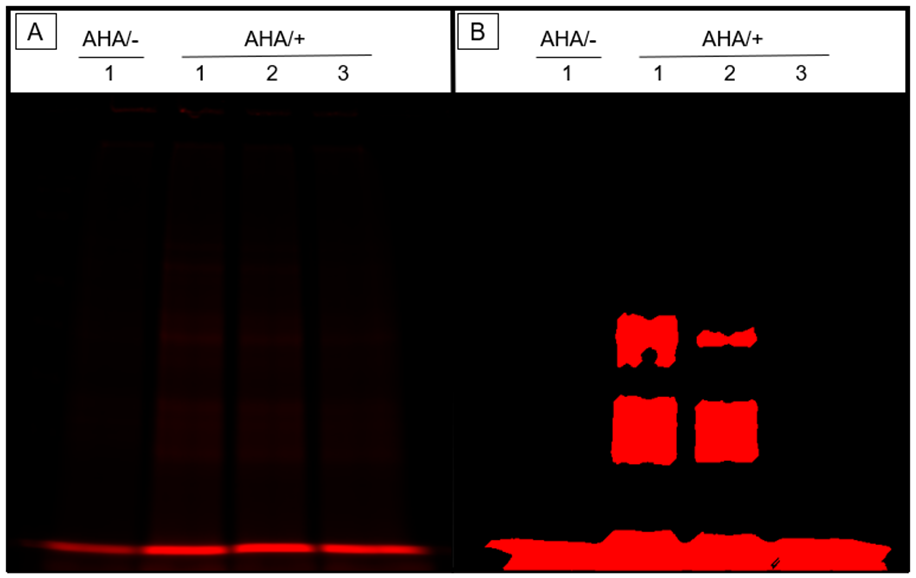

Supplementary Figure 5: SDS-PAGE to control the successful BONCAT labelling using click chemistry with fluorescent dye. Comparison of fluorescence intensity of (1-3) replicates incubated (AHA/+) with AHA and (1) replicate not incubated (AHA/-) with AHA of the biogas microbiome labelled with a fluorescent dye (Cyanine 5.5 DBCO) via click chemistry. 10 µg of the labelled proteins were loaded onto an SDS page followed by a fluorescence scan (Intensity: 0.5). A: original image; B: contrast and colour correction of the original image for better visibility of the bands.

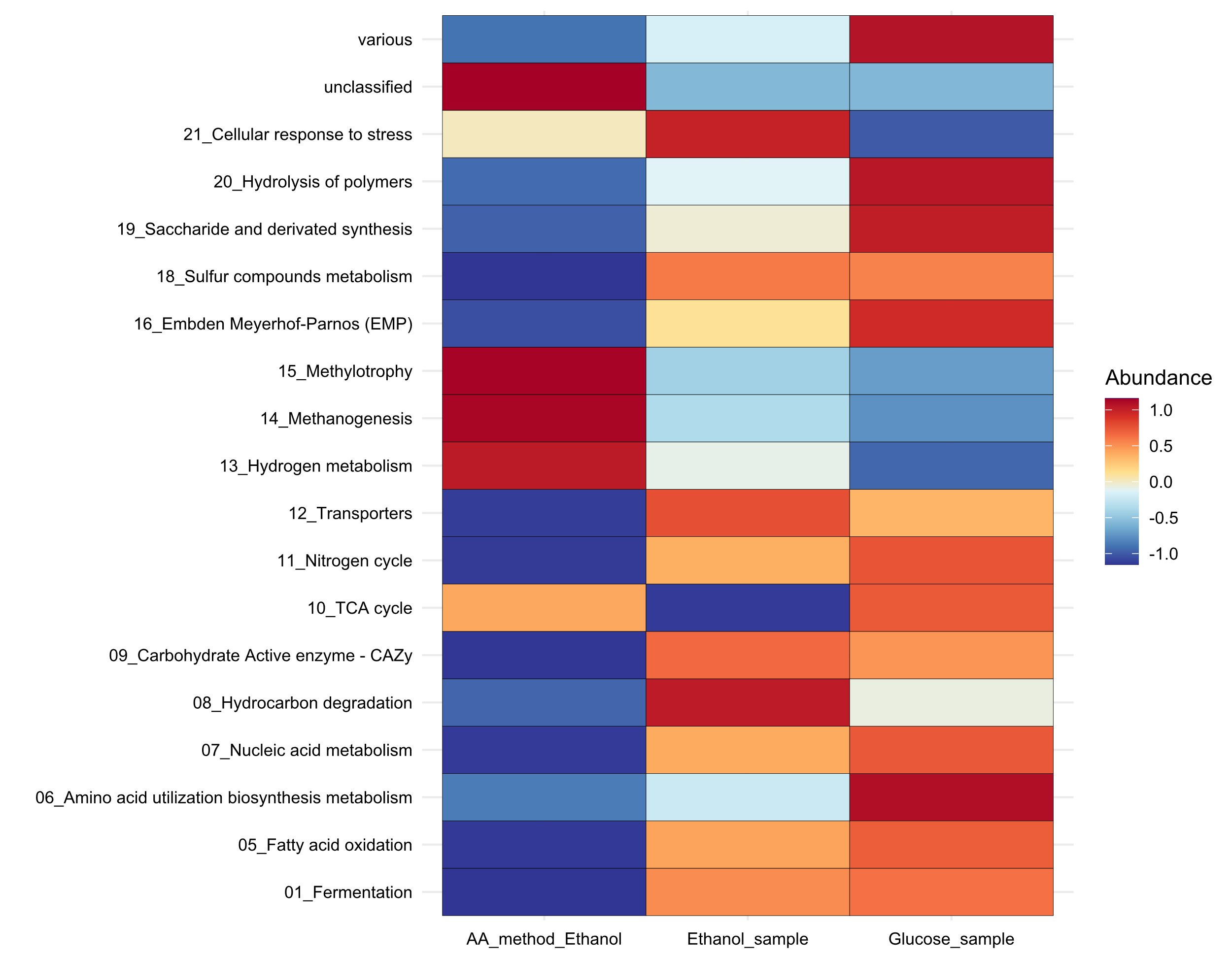

Supplementary Figure 6: The FOAM-based biological processes were analyzed using a heatmap created with R Studio (2023.06.0 Build 421) and ggplot2 (3.5.0). The data were normalized using a z-score per row, based on the average relative spectral count of two replicates. AA_method_Ethnaol: Enriched ethanol sample using the AA method, Ethanol_sample: Ethanol sample without enrichment, Glucose_sample: Glucose sample without enrichment See also Supplementary Table 5.

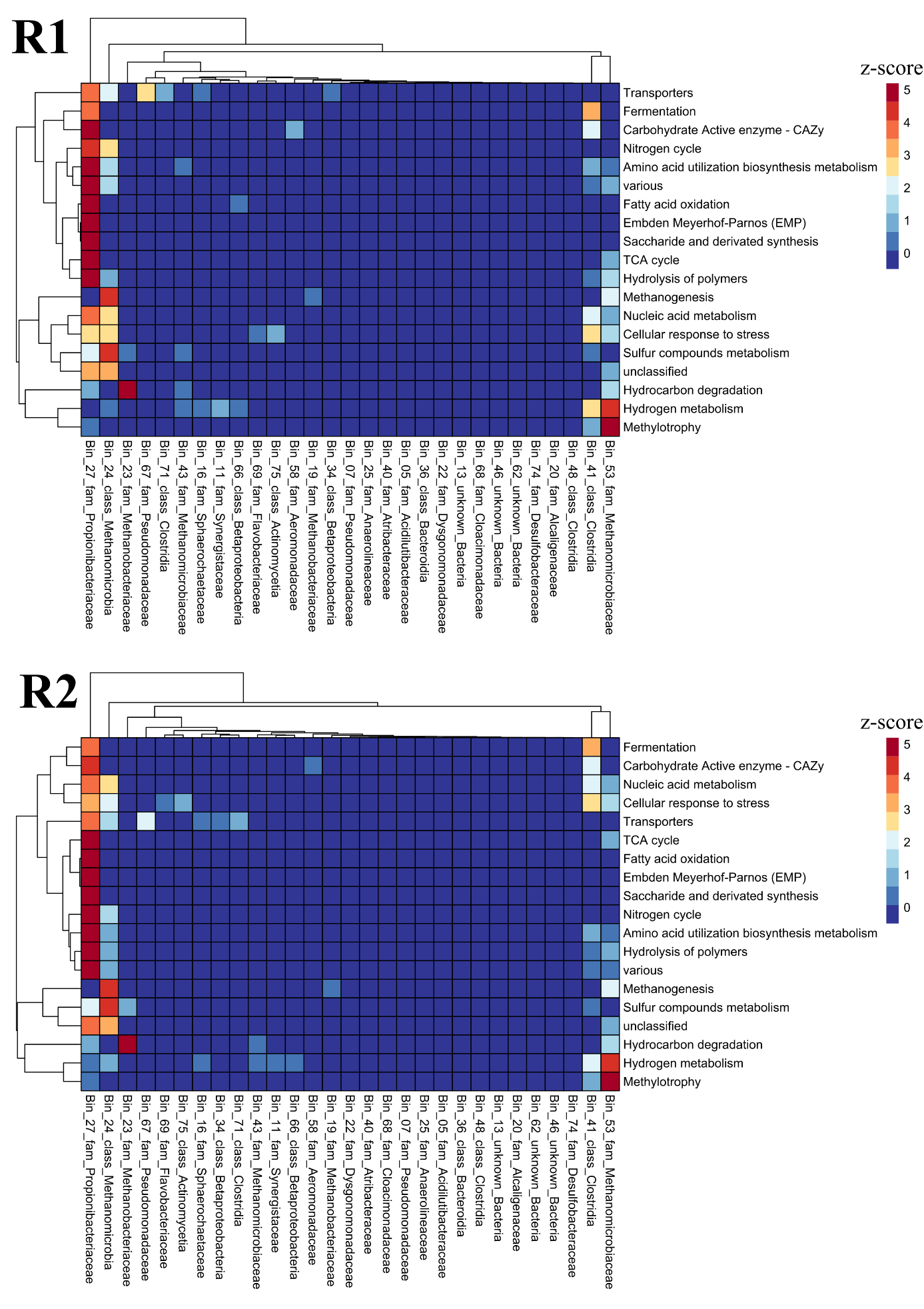

Supplementary Figure 7: Alternated biological processes in the MC due to ethanol addition. The functional ontology assignments for metagenomes of all proteins and MAGs without enrichment were analysed using a heatmap created with R Studio (2023.06.0 Build 421) and pheatmap (1.0.12). Previously, the protein abundances were normalised by dividing the spectral count of each protein by the total spectral abundance of the sample. Afterwards, a pivot table were generated to sum up the relative abundances of the proteins based on the functional assignment and the MAG. The functional ontology assignments were normalized using a z-score per row (over all MAGs). See also Supplementary Table 5

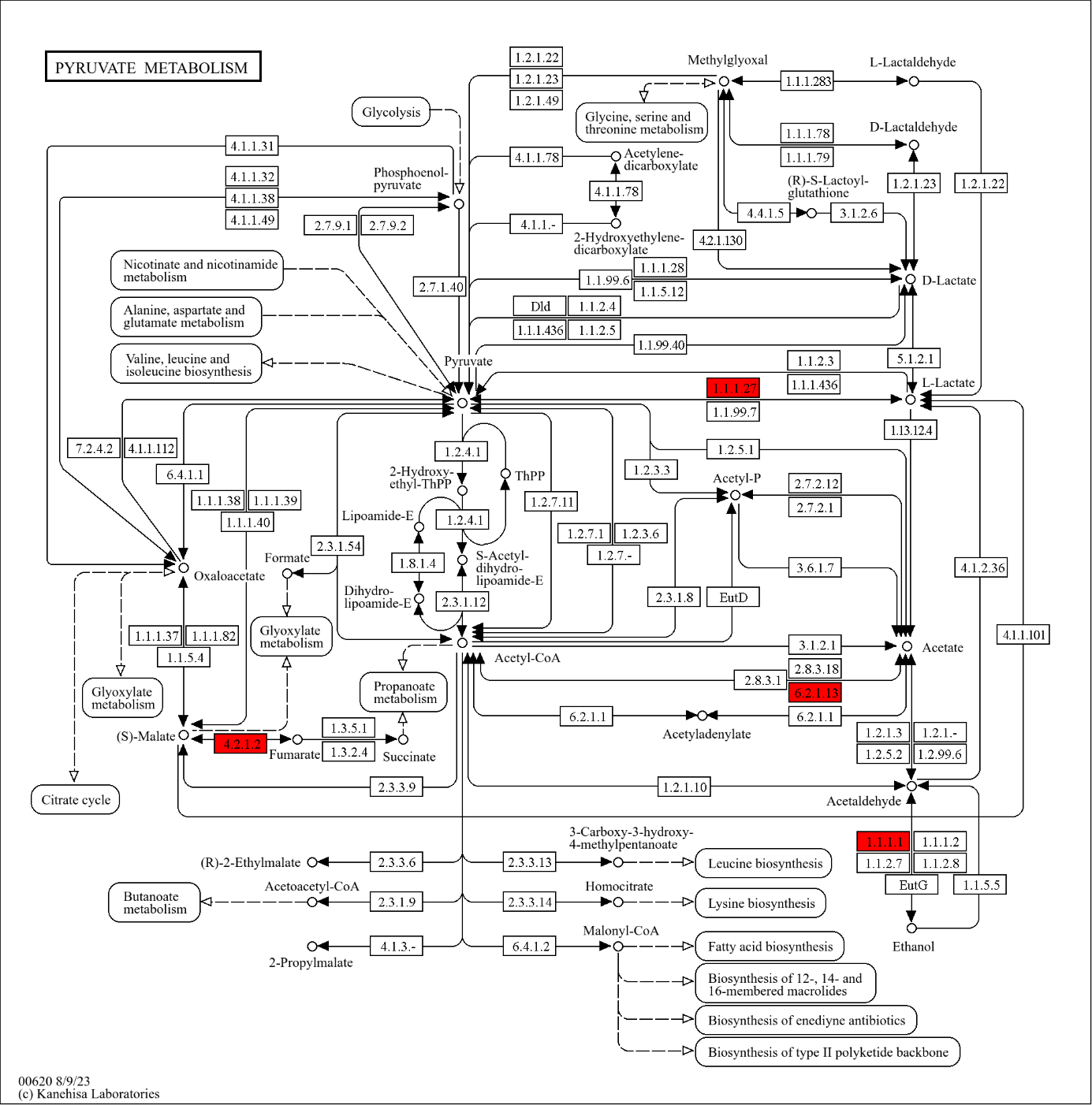

Supplementary Figure 8: KEGG map of all identified proteins from pyruvate metabolism in Bin41 (map 00620)

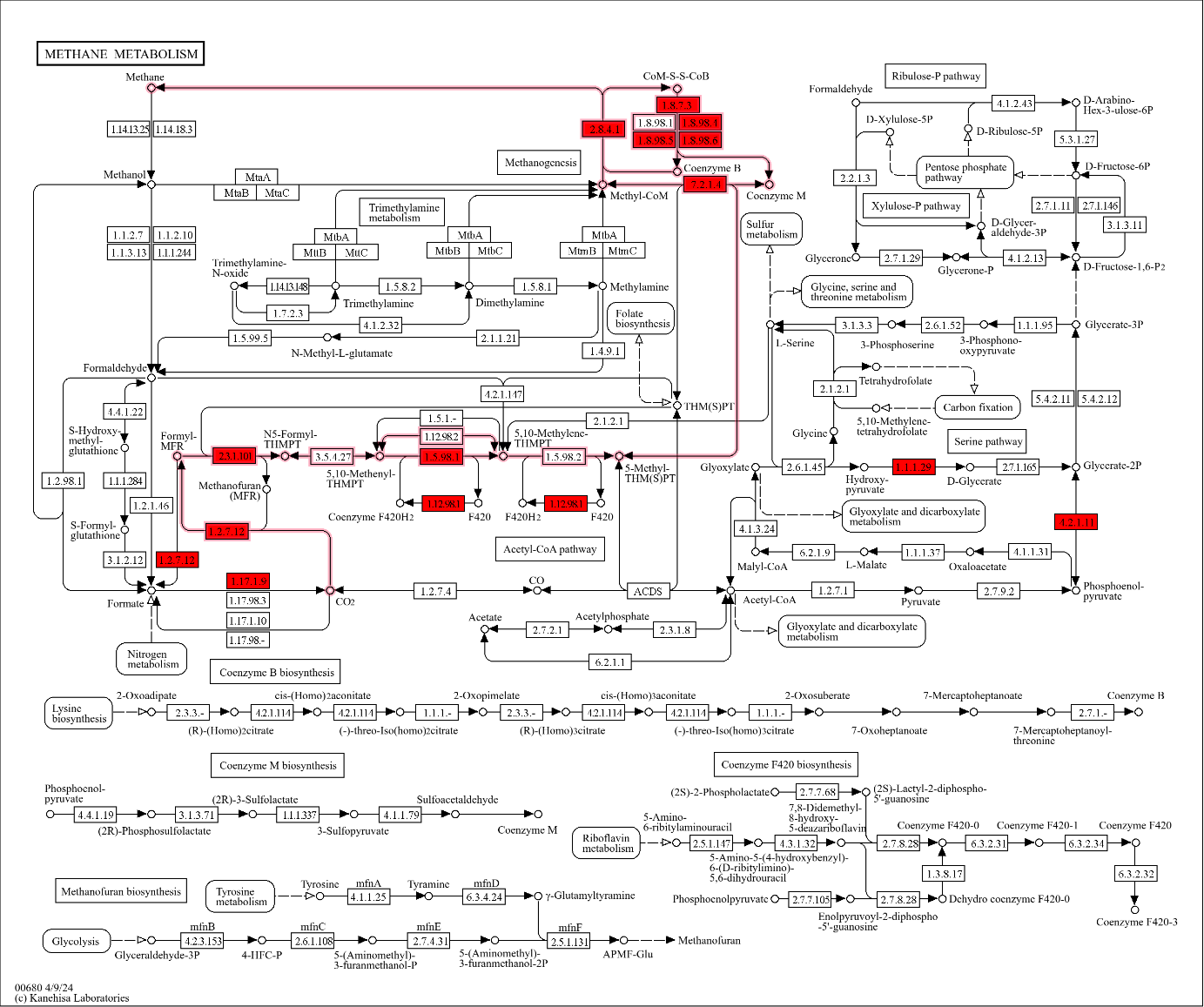

Supplementary Figure 9: KEGG map of all identified proteins from methane metabolism (map 00680) from Bin53

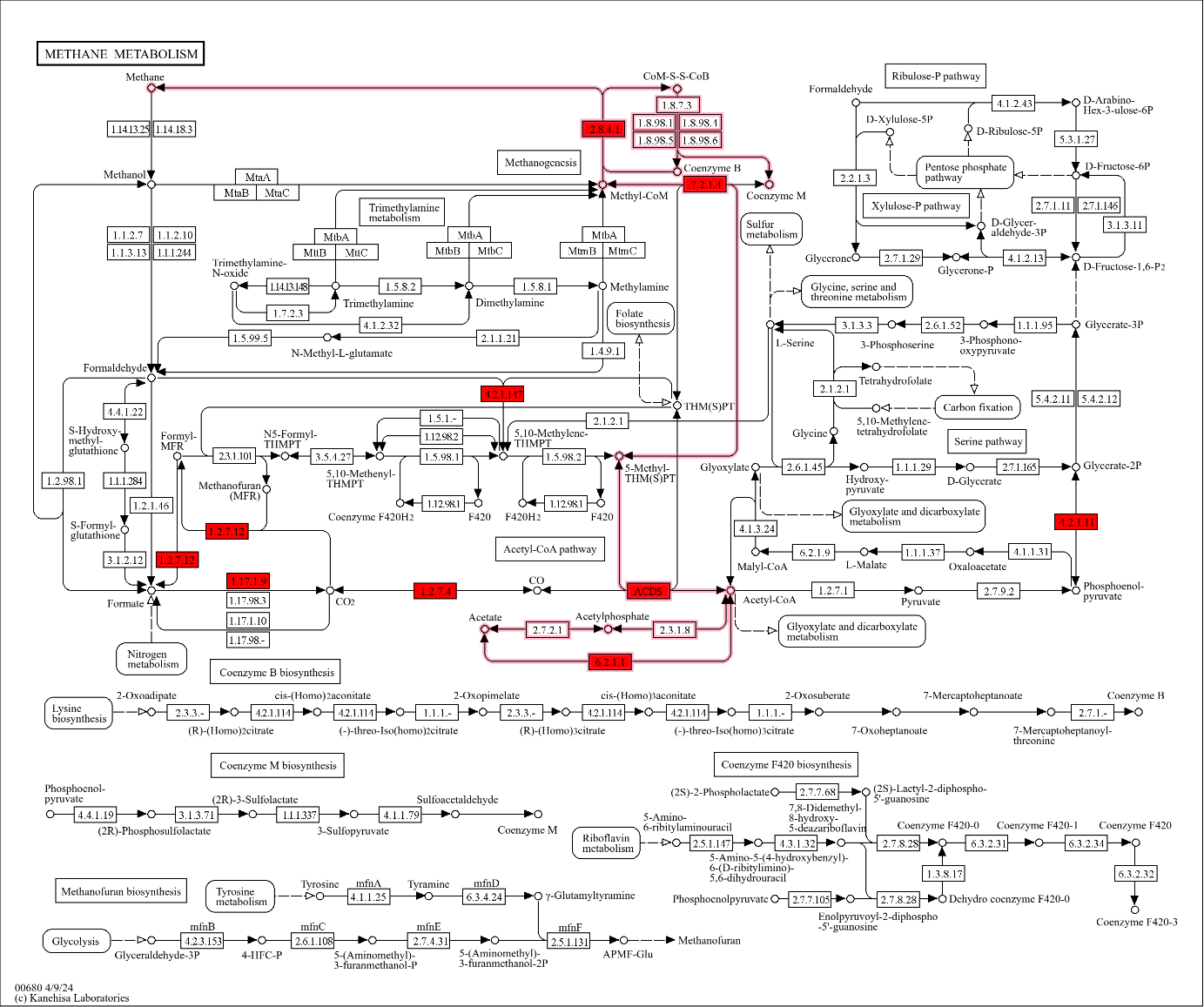

Supplementary Figure 10: KEGG map of all identified proteins from methane metabolism (map 00680) from Bin24

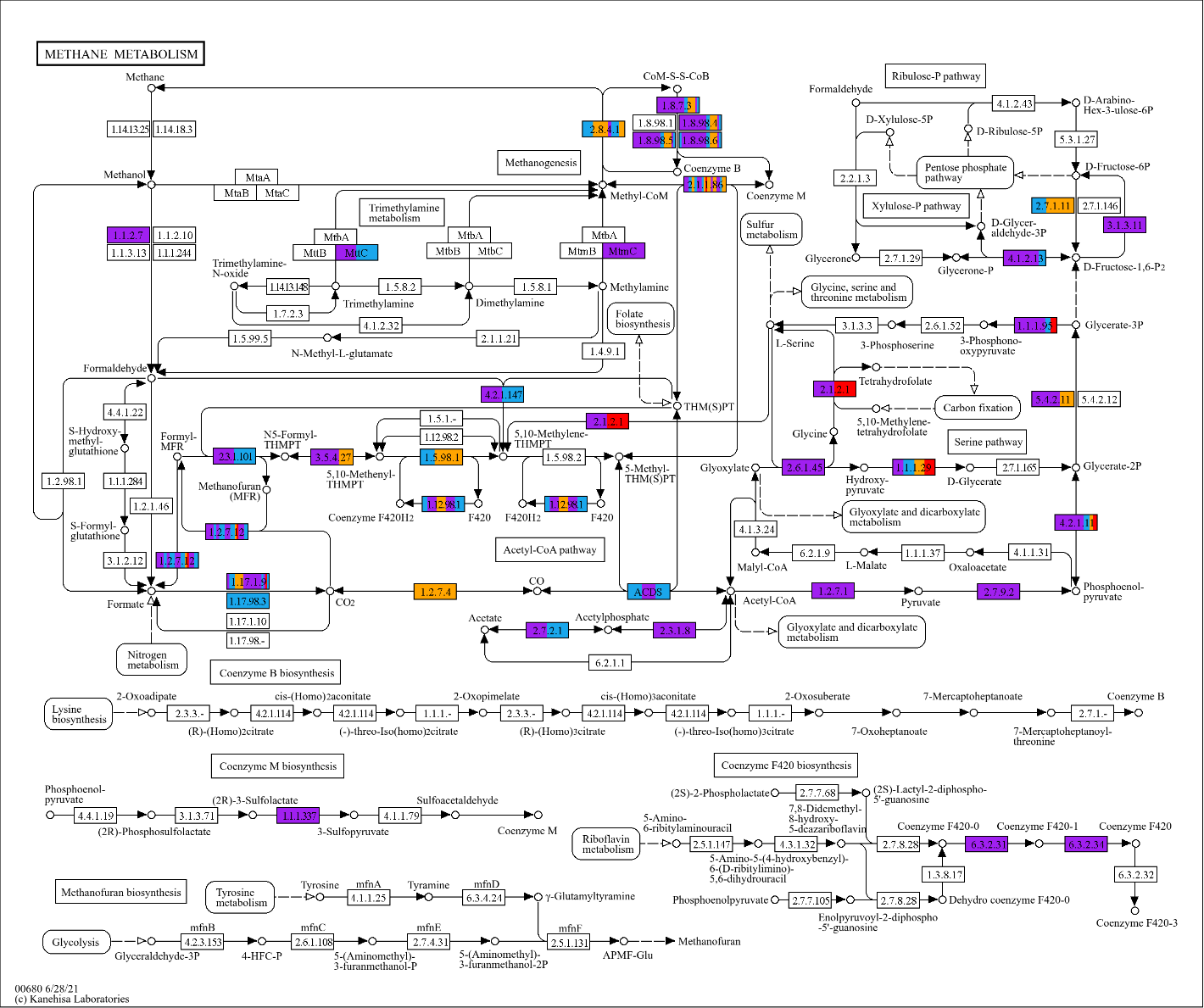

Supplementary Figure 11: KEGG map of all identified proteins from methane metabolism (map 00680). **Red** indicates the metaproteins unique identified with the developed AA method. **Orange** indicates the metaproteins enriched at least 2-fold with the developed AA method. **Purple** represents all metaproteins unique identified in the not enriched sample. **Blue** shows the metaproteins identified with the AA method and the not enriched sample. Boxes with several colours show that there are several similar proteins with different abundances for this step. Blank boxes were not identified in any sample. Samples were normalized by dividing the number of spectra of each metaprotein by the sum of the spectra of the sample. The abundance of each protein was averaged over 2 replicates, and the relative abundance ratio between the untreated sample and elution was calculated to compare the enrichment [6–8].

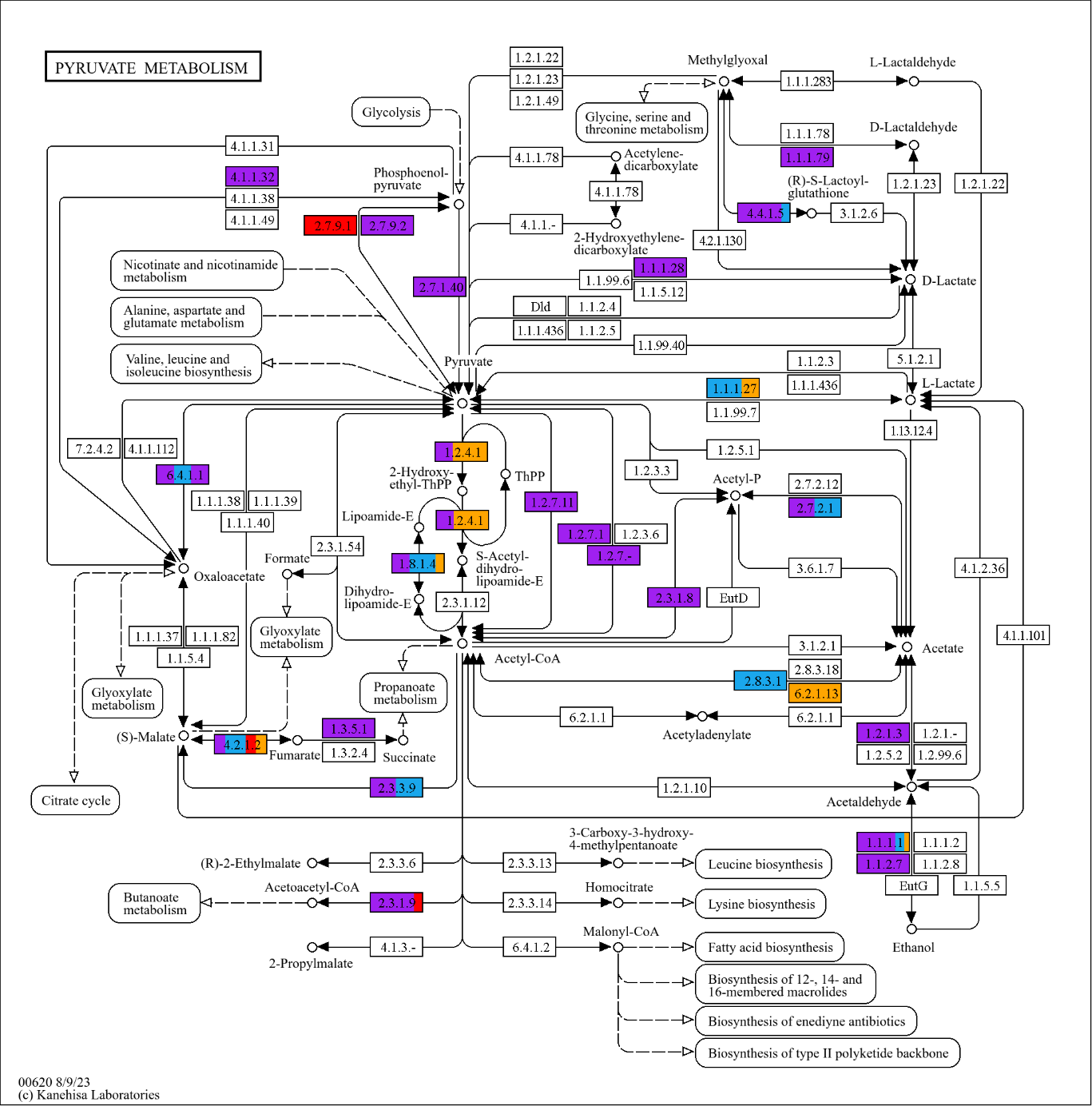

Supplementary Figure 12: KEGG map of all identified proteins from pyruvate metabolism (map 00620). **Red** indicates the metaproteins unique identified with the developed AA method. **Orange** indicates the metaproteins enriched at least 2-fold with the developed AA method. **Purple** represents all metaproteins unique identified in the not enriched sample. **Blue** shows the metaproteins identified with the AA method and the not enriched sample. Boxes with several colours show that there are several similar metaproteins with different abundances for this step. Blank boxes were not identified in any sample. Samples were normalized by dividing the number of spectra of each metaprotein by the sum of the spectra of the sample. The abundance of each protein was averaged over 2 replicates, and the relative abundance ratio between the untreated sample and elution was calculated to compare the enrichment [6–8].
